# *SNCA* triplication shapes neuronal extracellular vesicle biology and promotes microglial activation in patient-derived iPSC-based models of Parkinson’s disease

**DOI:** 10.64898/2026.09.09.750343

**Authors:** Ioanna Boumpoureka, Johanna Heider, Sylvie Delcambre, Felix B. Kleine Borgmann, Arnaud Poret, Ibrahim Boussaad, Johannes H. Wilbertz, Peter Sommer, Michel Mittelbronn, Anne Grunewald, Vyron Gorgogietas, Rejko Kruger

**Affiliations:** Luxembourg Centre for Systems Biomedicine (LCSB), Esch-sur-Alzette, Luxembourg; Ksilink, Strasbourg, France; Centre Hospitalier de Luxembourg (CHL), Luxembourg, Luxembourg; Transversal Translational Medicine, Luxembourg Institute of Health (LIH), Strassen, Luxembourg; Parkinson’s Research Clinic, Centre Hospitalier de Luxembourg (CHL), Luxembourg, Luxembourg; Luxembourg Centre of Neuropathology (LCNP), Dudelange, Luxembourg; Department of Life Science and Medicine (DLSM), University of Luxembourg, Esch-sur-Alzette, Luxembourg; Hôpitaux Robert Schuman, 2540 Luxembourg, Luxembourg; Division of Neuropathology, Department of Pathology and Neuropathology, Medical Faculty, University of Cologne, Germany; Mímirvion, Saarbrücken, Germany; Institut de Génétique et de Biologie Moléculaire et Cellulaire (IGBMC), University of Strasbourg, CNRS, Illkirch, France

**Author notes:** Corresponding author Correspondence to Vyron Gorgogietas, (,). These authors contributed equally and share senior authorship.

**Keywords:** Extracellular vesicles, Parkinson’s disease, stem cells, neurons, microglia

## Abstract

Extracellular vesicles (EVs) are emerging as key mediators of intercellular communication and potential biomarkers in Parkinson’s disease (PD), yet how disease-causing genetic alterations shape neuronal EV biology remains incompletely understood. Here, we used PD-patient iPSC-derived midbrain dopaminergic neurons (mDANs) harboring SNCA triplication (SNCA-4x), gene-corrected controls (SNCA-GC), and SNCA knockout (SNCA-KO) to investigate the impact of a-synuclein overexpression on neuronal-enriched EV (nEV) biology and neuron to microglia communication. SNCA-4x mDANs exhibited marked transcriptional alterations in pathways related to vesicle trafficking and extracellular matrix organization. Using an optimized isolation workflow, SNCA-4x neurons released significantly more and smaller nEVs enriched in a-synuclein, mitochondrial DNA (mtDNA), and PARK7/DJ-1 mRNA, while displaying reduced acetylcholinesterase activity. Functionally, SNCA-4x-derived nEVs were taken up more efficiently by isogenic control iPSC-derived microglia than SNCA-GC nEVs and induced stronger pro-inflammatory activation than both SNCA-GC nEVs and untreated microglia, characterized by altered microglial morphology and increased TNF-α and IL-1β, consistent with damage-associated molecular pattern (DAMP)-mediated signaling. Together, these findings demonstrate that SNCA-4x reshapes the properties and molecular cargo of nEVs, enhancing their capacity to trigger microglial activation, and identify EV-associated α-synuclein, mtDNA, and PARK7 mRNA as candidate mechanistic and biomarker features in PD.

**Graphical abstract:** *SNCA* triplication reshapes neuronal-enriched extracellular vesicle biology: compared with isogenic gene-corrected neurons, iPSC-derived SNCA-4x dopaminergic neurons released more and smaller nEVs carrying elevated levels of α-synuclein and mitochondrial DNA compared to the SNCA-GC dopaminergic neurons. These nEVs were taken up more efficiently by SNCA-GC iPSC-derived microglia and triggered pro-inflammatory activation, with increased levels of TNF-a and IL-1β compared to the SNCA-GC nEVs.

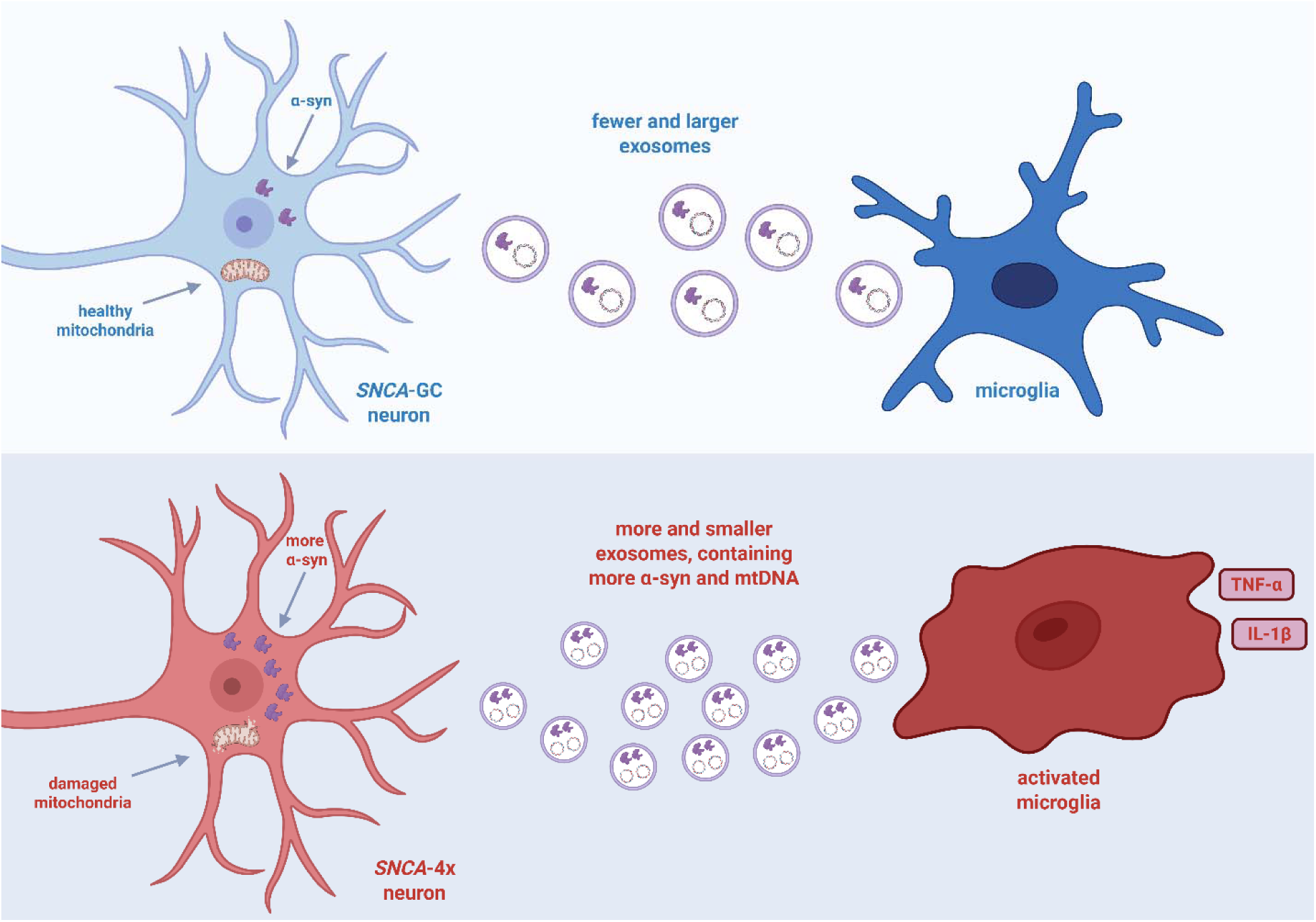

## 1. Introduction

### 1.1 Background

Parkinson’s disease (PD) is the second most prevalent neurodegenerative disorder worldwide, characterized clinically by progressive motor symptoms including bradykinesia, resting tremor, rigidity, and postural instability, along with non-motor features such as autonomic dysfunction, sleep disturbances, and neuropsychiatric symptoms (Kalia & Lang, 2015; Davie, 2008). The hallmark neuropathological feature of PD is the progressive loss of dopaminergic neurons in the substantia nigra pars compacta and the presence of Lewy bodies and Lewy neurites, which are inclusions predominantly composed of misfolded and aggregated α-synuclein (Kalia & Lang, 2015). At the molecular level, PD pathogenesis involves a complex interplay of mechanisms, including α-synuclein aggregation as the neuropathological hallmark contributing to neuronal toxicity (Mehra et al., 2019). Mitochondrial dysfunction is a well-established mechanism in both monogenic and idiopathic PD, characterized by decreased complex I activity, reduced mitochondrial membrane potential, increased ROS production, and altered mitochondrial morphology (Gao et al., 2022; Schapira et al., 1990). Furthermore, neuroinflammation plays a pivotal role in PD pathogenesis and progression, as evidenced by microglial activation and the release of inflammatory factors that directly damage neurons and exacerbate neuronal injury (Isik et al., 2023; Pajares et al., 2020).

Despite these established mechanisms of PD pathophysiology, a critical gap remains in our understanding of how PD pathology spreads throughout the brain. The mechanism by which a-synuclein spreads among cells and propagates pathological conversion from cell-to-cell remains largely unmapped. While prion-like propagation of misfolded α-synuclein has been proposed to underlie CNS proliferation of Lewy pathology, the specific mechanisms underlying cell-to-cell communication and the spread of PD pathology are not yet elucidated (Wu & Schekman, 2024; Valdinocci et al., 2017)

### 1.2 Study rationale

EVs are small (~30—200 nm), single-membrane, secreted vesicles that are enriched in selected proteins, lipids, nucleic acids, and metabolites (Chen et al., 2024). Their biogenesis involves the formation of the vesicles within multivesicular bodies (late endosomes) via the ESCRT pathway, and release occurs upon exocytic fusion of these organelles with the plasma membrane (Colombo et al., 2013). EVs function as crucial mediators of intercellular communication, transmitting signaling molecules and biomolecules between cells and playing important roles in development, immunity, tissue homeostasis, cancer, and neurodegenerative diseases (Kalluri & LeBleu, 2020).

EVs are increasingly recognized across multiple applications in neurodegenerative diseases: (1) understanding mechanisms of action of diseases by revealing how pathological biomolecules are transmitted among cells, (2) diagnostics, as EVs serve as biomarker sources for liquid biopsies from all human biofluids, reflecting pathological conditions and monitoring disease progression, and (3) therapeutics, as EVs are endogenous vesicles useful for drug delivery with biocompatibility, blood-brain barrier penetrability, metabolic stability, and target specificity, having been successfully loaded with therapeutic cargoes in PD models, including catalase protein (Haney et al., 2015), dopamine (Qu et al., 2018), catalase mRNA delivered by implanted designer-exosome-producing cells (Kojima et al., 2018), and α-synuclein-targeting siRNA delivered in rabies virus glycoprotein (RVG)-modified exosomes, which reduced α-synuclein mRNA, protein, and intraneuronal aggregates in the substantia nigra of transgenic mice (Cooper et al., 2014).

In PD, accumulating evidence suggests that EVs play an important role in disease progression. EVs may contribute to PD progression by facilitating the spread of pathological α-synuclein or by activating immune cells (Pinnell et al.,2021; Upadhya & Shetty, 2021). Extracellular a-synuclein is associated with EVs in vivo and in vitro, and the release of nEVs may represent a potential mechanism for propagation of pathological α-synuclein throughout the brain (Danzer et al., 2012; Xia et al., 2019; Alvarez-Erviti et al.,2011a). Microglial EVs have been confirmed to contain α-synuclein oligomers in the CSF of PD patients and can induce α-synuclein aggregation in neurons, supporting their role in disease progression (Guo et al., 2020). Beyond acting as vehicles for pathological protein, nEVs are increasingly recognized as regulators of microglial activity, with the direction of this signaling depending on the cargo carried by the donor neuron. EVs released by healthy neurons suppress microglial activation and reduce pro-inflammatory cytokine release (Peng et al., 2021), whereas EVs carrying pathological cargo activate microglia and propagate α-synuclein pathology (Xia et al., 2019; Guo et al., 2020). Both α-synuclein and mitochondrial DNA (mtDNA) are established damage-associated molecular patterns (DAMPs) capable of triggering innate immune responses and neuroinflammation (Zhang et al., 2010; Kim et al., 2013; West et al., 2015; Deus et al., 2022), providing a rationale for testing whether nEVs from α-synuclein-overexpressing neurons modulate microglial activation. In parallel, enzymatic activities carried by EVs have been proposed as disease-related readouts: acetylcholinesterase (AChE) activity is reduced in plasma EVs from PD patients and correlates negatively with disease severity (Shim et al., 2021), although AChE is not a generic EV marker and its activity differs between EV subpopulations (Liao et al., 2019).

### 1.3 Objective/hypothesis

Based on the accumulated evidence described above, which identifies α-synuclein as a critical regulator of EV-associated pathways, we wanted to investigate potential alterations in EV cargo composition. To this end, we modeled PD by using patient-derived iPSCs harboring a triplication of *SNCA*, a well-established monogenic model for PD. This approach enables us to evaluate the mechanisms underlying the propagation of disease-associated cargo within EVs, thereby providing insights into the putative cell-to-cell transmission of the α-synuclein phenotype characteristic for PD pathogenesis. Moreover, we aimed to identify novel molecular cargos with potential utility as future diagnostic biomarkers for clinical applications. Furthermore, we mechanistically interrogated the functional consequences of nEVs with altered cargo upon recipient cell types, with particular emphasis on microglia, which are the most physiologically relevant target in PD neuroinflammation. We aim to validate and characterize the mechanistic contributions of nEVs to PD pathophysiology, thereby advancing our understanding of neuronal EV-mediated intercellular communication in neurodegenerative disease progression.

## 2. Materials and Methods

### 2.1 Origin of iPSC lines and differentiation into mDA neurons and microglia

The *SNCA* iPSC lines used here, namely AST23 (SNCA-4x), AST23-2KO (SNCA-GC), and AST23-4KO (SNCA-KO), were a gift from Dr. Tilo Kunath (University of Edinburgh), who generated them. Devine et al. (2011) originally reported the parental AST23 line, and the derivation of the AST23-2KO and AST23-4KO clones was reported in Chen et al. (2019). We maintained iPSCs on Geltrex-coated dishes (Thermo Fisher Scientific) in mTeSR Plus (StemCell Technologies), refreshing the medium every second day and passaging weekly as clumps with 0.5 mM EDTA in DPBS (1:10 split; Thermo Fisher Scientific). Cultures were screened for mycoplasma every two weeks. The donor of the parental line gave written informed consent for inclusion of their data, was informed that they could not be identified from any published material, and all data were fully anonymised.

For dopaminergic differentiation, we applied a protocol adapted from Reinhardt et al. (2013). iPSCs were transferred into N2B27 base medium (49% DMEM/F12, #21331020; 49% Neurobasal, #21103049; 1:100 B27 without vitamin A, #12587010; 1:200 N2, #17502001; 1% (v/v) Glutamax, #35050061; and 1% (v/v) P/S, all from Thermo Fisher Scientific), to which we added 10 |jM SB-431542 (R&D Systems, #1614/10), 1 |jM Dorsomorphin (R&D Systems, #3093), 3 μM CHIR 99021 (CHIR; Axon Medchem, #2435) and 0.5 μM purmorphamine (PMA; Sigma, #SML0868). SB-431542 and Dorsomorphin were withdrawn after day 4, and cells were then held in N2B27 with CHIR, PMA and 150 μM ascorbic acid (AA; Sigma, #A4403). The neuroepithelium that formed by day 5 was manually excised, gently triturated, and replated onto Geltrex-coated wells; the resulting small-molecule neural progenitors (smNPCs) were expanded in N2B27 with CHIR, PMA and AA. For patterning, progenitors were plated on Geltrex-coated 6-well plates (3x10D cells/well) and cultured for 8 days in N2B27 containing 1 |jM PMA (Sigma-Aldrich, SML0868-25mg), 200 μM AA and 100 ng/ml FGF8b (Peprotech, 10-25). FGF8b was then removed between days 8 and 10 and PMA lowered to 0.5 μM; from day 10, cells were switched to maturation medium. Of note, N2B27 is serum-free and prepared without added EVs.

Microglia were generated by an established route (van Wilgenburg et al., 2013; Haenseler et al., 2017). Embryoid bodies (EBs) were first formed from iPSCs in mTeSR Plus (STEMCELL Technologies) containing 50 ng/ml BMP-4 (Invitrogen), 50 ng/ml VEGF (Invitrogen) and 20 ng/ml SCF (Miltenyi). EBs were moved to a low-attachment 6-well plate on day 4 with fresh EB medium, and on day 7 the medium was exchanged for X-VIVO 15 (Lonza) containing 25 ng/ml IL-3 (Invitrogen), 100 ng/ml M-CSF (Invitrogen), 2 mM Glutamax (Gibco), 1% P/S (Gibco) and 0.055 mM [3-mercaptoethanol (Gibco); at this point EBs were seeded into T75 “factory” flasks that continuously shed macrophage precursors. These factories were sustained for up to 6 months, with precursors collected periodically. Precursors were driven to mature microglia in advanced DMEM/F12 supplemented with N2, Glutamax, P/S, [3-mercaptoethanol, 100 ng/ml IL-34 (Peprotech) and 10 ng/ml GM-CSF (Peprotech). All cultures were kept at 37 °C and 5% CO_2_.

### 2.2 EV isolation

The neuronal media from mDANs cultured for 72 h were pooled together for EV isolation. The cellular debris was removed by centrifugation at 1000g for 10 min, followed by centrifugation at 15,000g for 40 min to remove microvesicles. The supernatant was filtered using a glass filtration system of 0.22 microns. For sucrose-based isolation, neuronal media were loaded slowly over 3 mL of 30% sucrose solution (prepared in 1 × phosphate-buffered saline), forming a layer, and centrifuged at 100,000g, 4 °C for 90 min using Optima MAX-TL ultracentrifuge in swinging Bucket rotor (Beckman, Coulter). Then the supernatant was discarded, and the sucrose layer (∼3 mL) was resuspended in 1 × DPBS and ultracentrifuged at 100,000g at 4 °C for 90 min to pellet down the EVs. After this, the EVs were resuspended in 500 pL 1 x DPBS and stored at - 80 °C for further use, in case of transmission electron microscopy or nanoparticle tracking analysis. For all the other downstream applications, EVs were precipitated using an equal volume of precipitation solution-16% polyethylene glycol with Mn (number average molecular weight) of 6000 (Sigma, 81260) in combination with DPBSx1 and sodium chloride (1DM) to make a two-fold concentrated (2x) stock solution, shaking at 4 °C overnight. The next day, the EVs were centrifuged at 15,000g at 4 °C, and the EV pellet could be stored at -80°C for further use.

### 2.3 EV characterization

For electron microscopy, samples diluted to several concentrations in ddH_2_O were applied (1 μ L per grid) to formvar/carbon-coated copper grids (ultra-thin, 200 mesh; EMS 215-412-8400) that had been activated in a Cressington 208 glow-discharge unit. Grids were rinsed three times in ddH_2_O and negatively stained with uranyl acetate. Images were collected on a Gemini SEM 300 (Zeiss) operated at 30 kV in sTEM mode.

Particle concentration and hydrodynamic size distributions were determined by nanoparticle tracking on a NanoSight NS300 (Malvern Instruments, UK) with NTA software v3.2. After priming the instrument with PBS (pH 7.4) and equilibrating to 25 °C, tracking accuracy was confirmed using 100 nm and 200 nm polystyrene bead standards (Malvern Instruments). We set the camera level to 14, at which all particles were detectable without saturation, and fixed the detection threshold at 5 to capture the majority of particles, while omitting ambiguous signals. Each sample was brought to 1 mL in PBS and diluted to roughly 100 particles/frame, and three 60 s videos were captured per sample. Mean size, mode (the dominant size population), and particle concentration (particles/mL) were extracted from the output.

EV protein markers were assessed by Western blotting. EV and cell pellets were lysed in RIPA buffer and total protein determined with the Pierce BCA kit (Thermo Fisher Scientific) per the manufacturer’s protocol. Thirty micrograms of EV lysate was resolved on 8 to 12% Bis-Tris gels (Thermo Fisher Scientific) by SDS-PAGE and transferred to nitrocellulose. Membranes were blocked for 1 h at room temperature in 5% (w/v) non-fat milk in TBS-T (10 mM Tris-HCl pH 7.5, 100 mM NaCl, 0.1% Tween 20), then probed with the primary antibodies listed in **Table 1**. Signals were revealed with the corresponding secondary antibodies (**Table 1**) and ECL reagent (Amersham, RPN2235) on an Odyssey XF imager (Li-Cor), and band intensities were quantified in ImageJ (RRID: SCR_003070).

**Table 1.** xxxx

| Antibodies | Dilution | Application | Cat# and RRID |
| --- | --- | --- | --- |
| OCT4 | 1:400 | ICC | Abcam, Cat# ab19857, RRID:AB_445175 |
| SOX2 | 1:100 | ICC | R&D Systems, Cat# AF2018, RRID:AB_355110 |
| Nestin | 1:1000 | ICC | R&D Systems, Cat# MAB1259, RRID:AB_2251304 |
| PAX6 | 1:100 | ICC | Covance, Cat# PRB-278P, RRID:AB_291612 |
| MAP2 | 1:1000 | ICC | Abcam, Cat# ab92434, RRID:AB_2138147 |
| TH | 1:500 | ICC | Sigma-Aldrich, Cat# AB152, RRID:AB_390204 |
| P2RY12 | 1:250 | ICC | Atlas Antibodies, Cat# HPA014518, RRID:AB_2669027 |
| TMEM119 | 1:250 | ICC | Sigma-Aldrich, Cat# HPA051870, RRID:AB_2681645 |
| α-synuclein | 1:1000 | ELISA | Abcam, Cat# ab138501, RRID:AB_2537217 |
| β-actin | 1:1000 | WB | Cell Signaling, Cat# 4967, RRID:AB_330288 |
| α-synuclein | 1:1000 | WB/ELISA | BD Biosciences, Cat# 610787, RRID:AB_398108 |
| NCAM | 1:1000 | WB | Abcam, Cat# ab118291, RRID:AB_10899970 |
| CD9 | 1:1000 | WB | Cell Signaling, Cat# 13403S, RRID:AB_2732848 |
| Flotillin-1 | 1:1000 | WB | Abcam, Cat# ab41927, RRID:AB_941621 |
| CD63 | 1:1000 | WB | Novus Biologicals, Cat# NB100-77913-, RRID:AB_1084027 |
| Calnexin | 1:1000 | WB | Cell Signaling, Cat# 2679, RRID:AB_2228381 |
| Goat Anti-Mouse IgG HRP | 1:10000 | WB | Invitrogen, Cat# A24524, RRID:AB_2535993 |
|  | 1:5000 | ELISA |  |
| Goat Anti-Rabbit IgG HRP | 1:10000 | WB | Invitrogen, Cat# A24537, RRID:AB_2536005 |
|  | 1:5000 | ELISA |  |

### 2.4 Cell and EVs staining, immunocytochemistry and imaging

EVs diluted in DPBS were stained by adding them on top of dye preparations as follows: for carboxyfluorescein succinimidyl ester (CFSE) staining (Biolegend, 423801), 100D|jL of EV-containing DPBS solution was pipetted onto 100D|jL of a 40D|jM CFSE solution, and incubated for 2 h at 37°C, unless otherwise indicated. Tubes were mixed by flicking every hour. The CFSE-stained EVs were added to centrifugal filters of 10 kDa (Amicon Ultra 0.5, Millipore) and spun to remove the excess CFSE dye and to concentrate the EV volume. The concentrated CFSE-stained EVs were resuspended in culture media, quantified, and added to iPSC-derived microglia for 24 h. The nEV dose was normalised before treatment: CFSE-stained nEVs were quantified by NTA, and an equal number of particles per recipient microglial cell was applied (1000 nEVs/ microglial cell) for SNCA-4x and SNCA-GC conditions, so the read-outs reflect per-particle differences in uptake and not the higher secretion by SNCA-4x neurons.

Both iPSC-derived microglia and neurons were used for immunocytochemistry analysis. Briefly, microglia or neurons were plated in black 96-well plates and were then fixed with 4% paraformaldehyde in PBS for 15min at RT, followed by permeabilization and blocking in 0.4% Triton-X 100 (Carl Roth) and 2% bovine serum albumin (Sigma-Aldrich) in PBS for 1h at RT. Subsequently, permeabilized cells were stained with primary antibodies in incubation buffer (0.1% Triton-X and 0.2% bovine serum albumin in PBS) overnight at 4°C, washed three times with PBS, and incubated for 2h at RT with secondary antibodies in incubation buffer. The primary and secondary antibodies that were used in our studies are listed in **Table 1** along with the nuclear stain Hoechst 33342 (Invitrogen). Images were acquired with the Zeiss Cell Observer Spinning Disk microscope (Carl Zeiss MicroImaging GmbH).

Phalloidin iFluor-555 staining (StemCell Technologies) was used for the morphometric analysis of microglial cells, according to the manufacturer’s instructions. Briefly, after EV treatment cells were fixed and permeabilized as described above and were stained with phalloidin in a concentration of 1:1000 in PBS for 1h. After three PBS washes, z-stacks were acquired with the Zeis Cell Observer microscope and images were analyzed using the ‘Microglia Morphometry’ plugin in Fiji (RRID: SCR_002285), as described elsewhere (Martinez et al., 2023). More specifically, the microglial cell-shape descriptors extracted defines as circularity = 4tt x area/perimeter^2 (1.0 = perfect circle) and solidity = area/convex-hull area (proportion of a convex outline that is filled); both increase as cells retract their ramified processes and adopt a rounded, amoeboid, activated morphology.

EV uptake by microglia was analyzed using the Imaris image analysis software (Oxford Instruments, RRID: SCR_007370). The acquired z-stack images were used for 3D rendering of a microglial surface mask, based on the phalloidin staining. Subsequently, the number of exosomes localized within the microglia 3D mask was quantified using the spot detection tool to ensure that only exosomes uptaken by microglia were considered for the analysis (see Supplementary Figure for an example image of masks generated in Imaris). A consistent fluorescence intensity threshold for exosome detection was applied in all conditions. A surface mask was also generated for Hoechst-positive nuclei, and the number of exosomes detected was normalized to the number of Hoechst-positive nuclei per image.

### 2.5 EV acetylcholinesterase activity assay

The evaluation of EV acetylcholinesterase (AChE) activity was performed as follows: 25 μL of the EV lysate was added to 100 μL of phosphate buffer, then 1.25 mM acetylthiocholine (Sigma, 01480) and 0.1 mM 5,5’-dithiobis-2-nitrobenzoic acid (Sigma, D8130) were added to reach a final volume of 200 μL. The reaction mixture was incubated in 96-well plates at 37°C, and the absorbance at 412 nm was monitored continuously at a Cytation 5 plate reader (BioTek). The enzymatic activity was determined based on the change in absorbance observed after 20 minutes of incubation.

### 2.6 RNA-seq Library Preparation and Sequencing

Total RNA was extracted from samples using standard protocols. The concentration and quality of the extracted RNA were assessed prior to library preparation. Library preparation was performed using 3-mRNA sequencing, selectively enriching for polyadenylated transcripts to capture messenger RNA. Sequencing was conducted by Life & Brain (Bonn, Germany), generating approximately 20 million raw reads per sample.

### 2.7 RNA-seq Data Processing

Raw sequencing reads underwent initial quality control and preprocessing using fastp, which performed read trimming and filtering to remove low-quality bases and adapter sequences. The quality of the processed reads was assessed using FastQC (RRID: SCR_014583) to ensure data integrity prior to alignment. Reads were aligned to the Homo sapiens GRCh38 reference genome obtained from Ensembl (RRID: SCR_002344) using STAR (RRID: SCR_004463) in a two-pass mode to optimize splicing detection. Gene annotations were provided via the corresponding Ensembl GTF file. Following alignment, Samtools (RRID: SCR_002105) was used to remove duplicate reads, thereby minimizing PCR amplification bias. Quantification was performed using HTSeq (RRID: SCR_005514), generating a count matrix representing the expression levels of genes across conditions.

### 2.8 Differential Expression Analysis

Gene expression comparisons were conducted between a test condition characterized by *SNCA* triplication (SNCA-4x) and an isogenic gene-corrected control (SNCA-GC) condition. The count matrix was filtered to retain only protein-coding genes before differential expression analysis. Differential expression was assessed using DESeq2 (RRID: SCR_000154), applying a fold-change threshold of 1.5 and a false discovery rate (FDR) cutoff of 0.05 to identify significantly altered genes.

### 2.9 Functional Enrichment Analysis

To gain biological insights, the fgsea R package was utilized for over-representation analysis (ORA) and gene set enrichment analysis (GSEA). ORA was conducted on differentially expressed genes that met the defined criteria, testing for enrichment in Gene Ontology (GO) terms and Human Phenotype Ontology (HPO) categories, with an FDR cutoff of 0.05. GSEA was subsequently performed using fgsea, ranking differentially expressed genes by FDR-adjusted significance and evaluating enrichment across the same gene sets (GO and HPO) with an FDR threshold of 0.05.

### 2.10 EV nucleic acid isolation, Real-Time PCR, and digital PCR

Based on the manufacturer’s recommendation, the EV pellets are resuspended in 1X DPBS and treated with Nuclease Mix (Cytiva, #80650142) to remove exogenous-non-EV nucleic acids. Then, using the TRIzol-based method, total RNA was obtained from the upper aqueous phase of TRIzol LS (Thermo Fisher Scientific, #10296010) and EV DNA from the lower organic phase. More specifically, the aqueous phase was further processed and isolated for RNA isolation using the RNeasy Plus Mini kit (Qiagen, #74134), following the manufacturer’s instructions. For the DNA isolation, the lower organic phase was precipitated by adding 1200 μl ethanol and 20 |jL sodium acetate, followed by centrifugation at 16,000 RPM for 30 minutes. The DNA pellet was further processed and isolated using the DNeasy Blood and Tissue Mini kit (Qiagen, #69504), following the manufacturer’s instructions.

The isolated EV RNA was converted to cDNA using the High-Capacity cDNA Reverse Transcription Kit (Applied Biosystems, #4368814), following the manufacturer’s instructions. Gene expression was assessed on a LightCycler 480 II (Roche Diagnostics), using the PowerTrack SYBR Green Mastermix (Thermo Fisher Scientific, A46012). The primer sets used are available in **Table 2**.

**Table 2.**
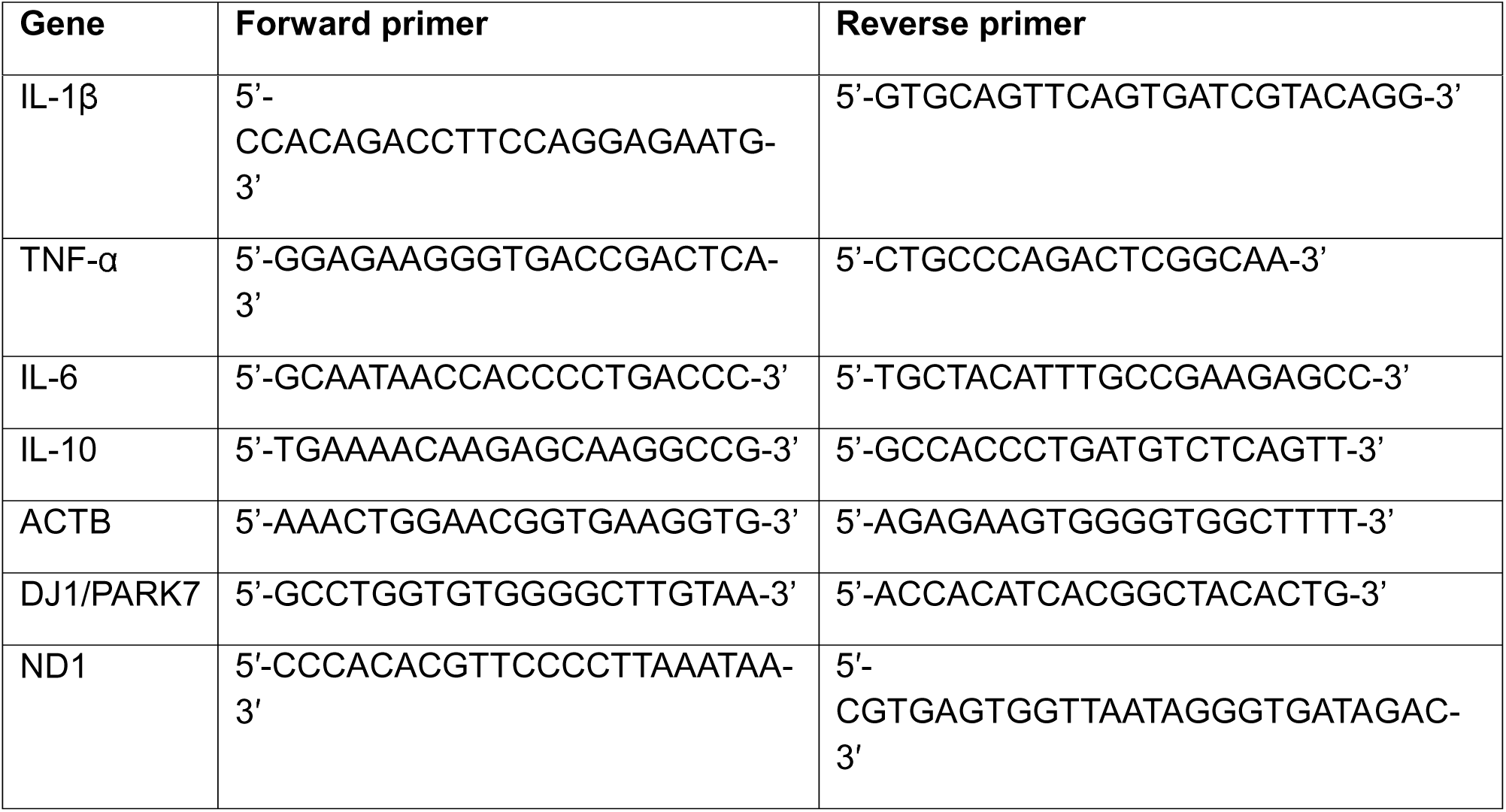
xxxx

We used isolated EV DNA to quantify mitochondrial DNA integrity and copy number by digital PCR (dPCR) on the QuantStudio Absolute Q Digital PCR System in the MAP16 format, following an established protocol (Ghelfi et al., 2025). Four targets were measured in a multiplexed hydrolysis-probe assay: the mitochondrial genes ND1 (VIC/MGB probe, Assay ID APAANDG, Cat #44485) and ND4 (ABY/QSY probe, custom, Article No. CCU002NR), the mitochondrial D-loop (JUN/QSY probe, custom, Article No. CCU002NR), and the single-copy nuclear gene B2M (FAM/MGB probe, Assay ID APCFGXE, Cat #43320). Assays were used at a final 1X concentration, with no more than two MGB-quenched assays combined per reaction to preserve fluorescence separation. Each reaction was assembled to a final volume of10 μL, containing 2 μL of 5X Absolute Q DNA Digital PCR Master Mix (Thermo Fisher Scientific, #A52490), 0.5 μL of each 20X probe, template DNA, and nuclease-free water (Thermo Fisher Scientific, #R0582), with one extra reaction prepared per run to account for pipetting loss. For each well, 9 μL of the sample and master-mix reaction was combined with 15 μL of isolation buffer on a MAP16 plate (QuantStudio Absolute Q MAP16 Plate Kit and Master Mix, Thermo Fisher Scientific, #A53301). Thermal cycling comprised a pre-heat at 96 °C for 10 min followed by 40 cycles of 96 °C for 5 s and 60 °C for 30 s. Runs were analysed with the Absolute Q software, requiring clean partition images and clear separation of positive and negative partitions. Mitochondrial DNA status was expressed as three ratios: ND1/B2M as an index of mtDNA copy number per cell, ND4/ND1 as an index of deletion load, and D-loop/ND1 as an index of mtDNA integrity and replication.

### 2.11 ELISA for a-synuclein and cytokines

EV-associated α-synuclein (predominantly monomeric) was quantified with a sensitive ELISA adapted from Emmanouilidou et al. (2011). Capture relied on the monoclonal Syn-1 antibody (BD Biosciences), which recognizes residues 15 to 123 of human α-synuclein, while detection used the MJFR1 antibody (Abcam, ab138501), which recognizes an epitope within the C-terminal region (residues ∼118-123) of full-length human α-synuclein. Corning Costar plates were coated overnight (24 h, RT) with Syn-1 at 0.5 μg/mL (50 μL/well) in 100 mM NaHCO_3_ (pH 9.3) and then stored at 4 °C for up to 2 to 3 weeks. Following three washes in wash buffer (50 mM Tris-HCl, 150 mM NaCl, 0.04% Tween-20), 50 μL of EV lysate or recombinant α-synuclein standard, diluted in TBST/BSA (10 mM Tris-Cl pH 7.6, 100 mM NaCl, 0.1% Tween-20, 1% BSA), was loaded and left to bind for 2.5 h at 37 °C. Wells were washed three times, incubated with 50 μ L detection antibody (in TBST/BSA) for 1 h at RT, washed again, and incubated with 50 μL HRP-conjugated secondary antibody (Invitrogen, A24537) for a further 1 h at RT. After a final wash, 100 μL HRP substrate (BioLegend, 421101) was added and developed for at least 15 min at RT; the reaction was terminated with 50 μL 1 M H_3_PO_4_ and absorbance read at 450 nm on a Cytation 5 reader.

Secreted cytokines (IL-1β, TNF-α and IL-10) were measured in microglial conditioned media using high-sensitivity kits: the Human IL-1β High Sensitivity ELISA (Thermo Fisher Scientific, BMS224-2HS), the LEGEND MAX Human TNF-α High Sensitivity ELISA (BioLegend, 430217), and the LEGEND MAX Human IL-10 High Sensitivity ELISA (BioLegend, 430617), each according to its manufacturer’s protocol. Microglia had been treated with SNCA-GC nEVs, SNCA-4x nEVs, LPS + IFN-γ (positive control), or vehicle (untreated control); supernatants were harvested, clarified by centrifugation, and assayed directly. In brief, samples and standards were bound on antibody-coated plates, followed by detection antibody and enzyme conjugate, substrate development, and a stop step, with absorbance recorded at 450 nm on a Cytation 5 reader. Concentrations were interpolated from recombinant-standard curves.

### 2.12 Software and statistical analysis

ImageJ was used for the image analysis and the downstream quantification of the selected features. Statistical analyses were performed using GraphPad Prism (version 9.5.1, RRID: SCR_002798). Unless otherwise specified, data were obtained from independent experiments or from independent differentiation batches of hiPSC-derived neurons. The number of biological samples and independent experimental replicates used for each analysis is indicated in the corresponding figure legends. For comparisons between two groups, a two-tailed unpaired t-test was applied. For analyses involving a single independent variable, one-way analysis of variance (ANOVA) followed by Tukey’s post hoc multiple comparisons test was used. For experiments with more than one independent variable, two-way ANOVA followed by Tukey’s post hoc multiple comparisons test was performed. The specific statistical tests used are indicated in the respective figure legends. A p-value < 0.05 was considered statistically significant (\**p < 0.05, **p < 0.01, \*\*\**p < 0.001, ****p < 0.0001). For the preparation of this work, the author used Grammarly and Claude AI to improve grammar, language quality, and readability. After using these tools, the author further reviewed and edited the content as needed and takes full responsibility for the content of the publication.

## 3. Results

### 3.1 Characterization of iPSC-derived midbrain dopaminergic neurons and microglia cells

To establish reliable cellular models for investigating SNCA-related pathology, we differentiated iPSCs harboring an *SNCA* triplication mutation (SNCA-4x), the respective isogenic control (SNCA-GC), and SNCA knockout (SNCA-KO) lines into midbrain dopaminergic neurons (mDANs) and microglia cells using established protocols (Fig. 1A). Stage-specific marker expression confirmed successful differentiation at each developmental stage: iPSCs expressed the pluripotency markers SOX2 and OCT4 (Fig. 1B), neural progenitor cells (NPCs) were positive for PAX6 and Nestin (Fig. 1C), differentiated mDANs expressed the pan-neuronal marker MAP2 and the dopaminergic marker tyrosine hydroxylase (TH) (Fig. 1D), and microglia cells displayed the microglial markers TMEM119 and P2RY12 (Fig. 1E). Western blot analysis confirmed that SNCA-4x mDANs produced significantly elevated levels of monomeric a-synuclein protein compared to isogenic control cells (Fig. 1F), validating the disease-relevant overexpression phenotype. An ELISA assay confirmed significantly elevated α-synuclein protein in SNCA-4x mDANs compared with both isogenic control and SNCA-KO cells (Fig. 1G).

**Figure 1.**
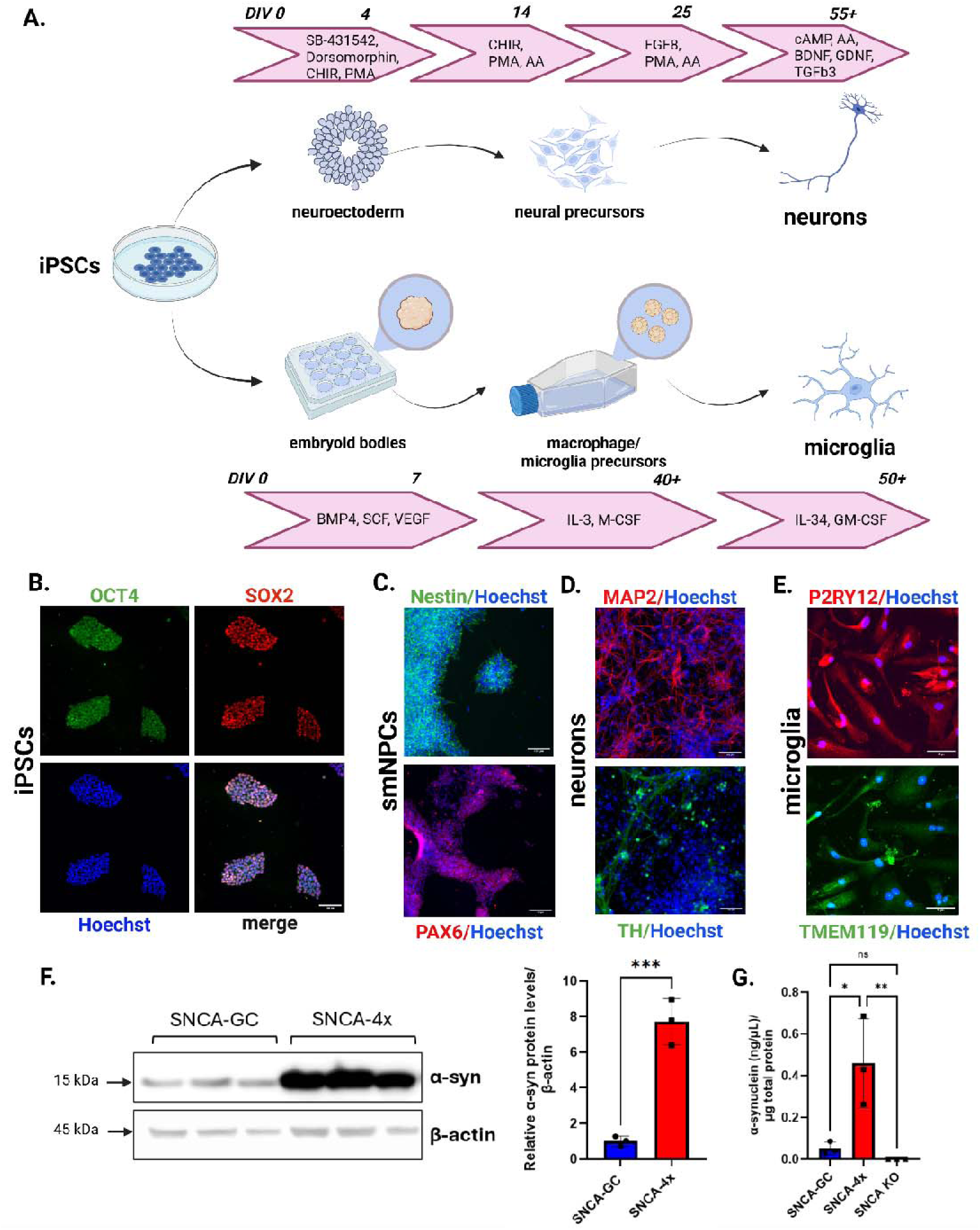
Generation and characterization of cellular models. (A) Schematic representation of the differentiation of iPSCs into either 1) neural progenitor cells (NPCs) and subsequently mature dopaminergic neurons (mDANs) or 2) macrophage-like microglia. (B) Immunocytochemical validation of iPSCs using the pluripotency markers OCT4 (green) and SOX2 (red). (C) Immunocytochemical validation of NPCs using Nestin (green) and PAX6 (red), and of (D) mDANs using MAP2 (red) and TH (green). (E) Immunocytochemical validation of microglia using the microglial markers P2RY12 (red) and TMEM119 (green). In all panels, nuclei are counterstained with Hoechst 33342 (blue). Scale bars: 100 μm for panels B and C, and 50 μm for panel D. (F) Western blot analysis of α-synuclein protein levels in day 30 mDANs derived from the SNCA-4x and SNCA-GC lines, followed by quantification of relative a-synuclein levels normalized to [3-actin in SNCA-4x (red) and SNCA-GC (blue) cells. (G) Absolute quantification of the protein levels of α-synuclein in cell lysates of SNCA-GC (blue), SNCA-4x (red) and SNCA-KO (purple). An unpaired t-test was performed using three biological replicates from independent experiments. ns, not significant; \**p < 0.05, **p < 0.01, \*\*\**p < 0.001.

### 3.2 SNCA-4x mDANs exhibit transcriptional alterations in extracellular vesicle-related pathways

To characterize molecular differences between SNCA-4x and SNCA-GC mDANs, and to determine how α-synuclein alters neuronal functions, we performed bulk RNA sequencing (RNA-seq) (Fig. 2). Principal component analysis (PCA) successfully distinguished SNCA-4x triplication samples from isogenic controls, demonstrating clear transcriptional divergence between genotypes (Fig. 2A). Differential expression (DE) analysis identified genes differentially expressed at a 5% false discovery rate (FDR), with top candidates including *GP2, PRSS23, CALCA, DSC2, LAMC2, MXRA5*, and *S100A6* (Fig. 2B). Gene ontology (GO) enrichment analysis revealed significant alterations in biological processes related to neuronal development, neuronal differentiation, and synaptic activity (Fig. 2C-D). Notably, we observed significant enrichment in pathways associated with EVs, membrane organization, and extracellular matrix dynamics, suggesting that α-synuclein overexpression profoundly affects vesicle trafficking, presynaptic membrane dynamics, and intercellular communication.

**Figure 2.**
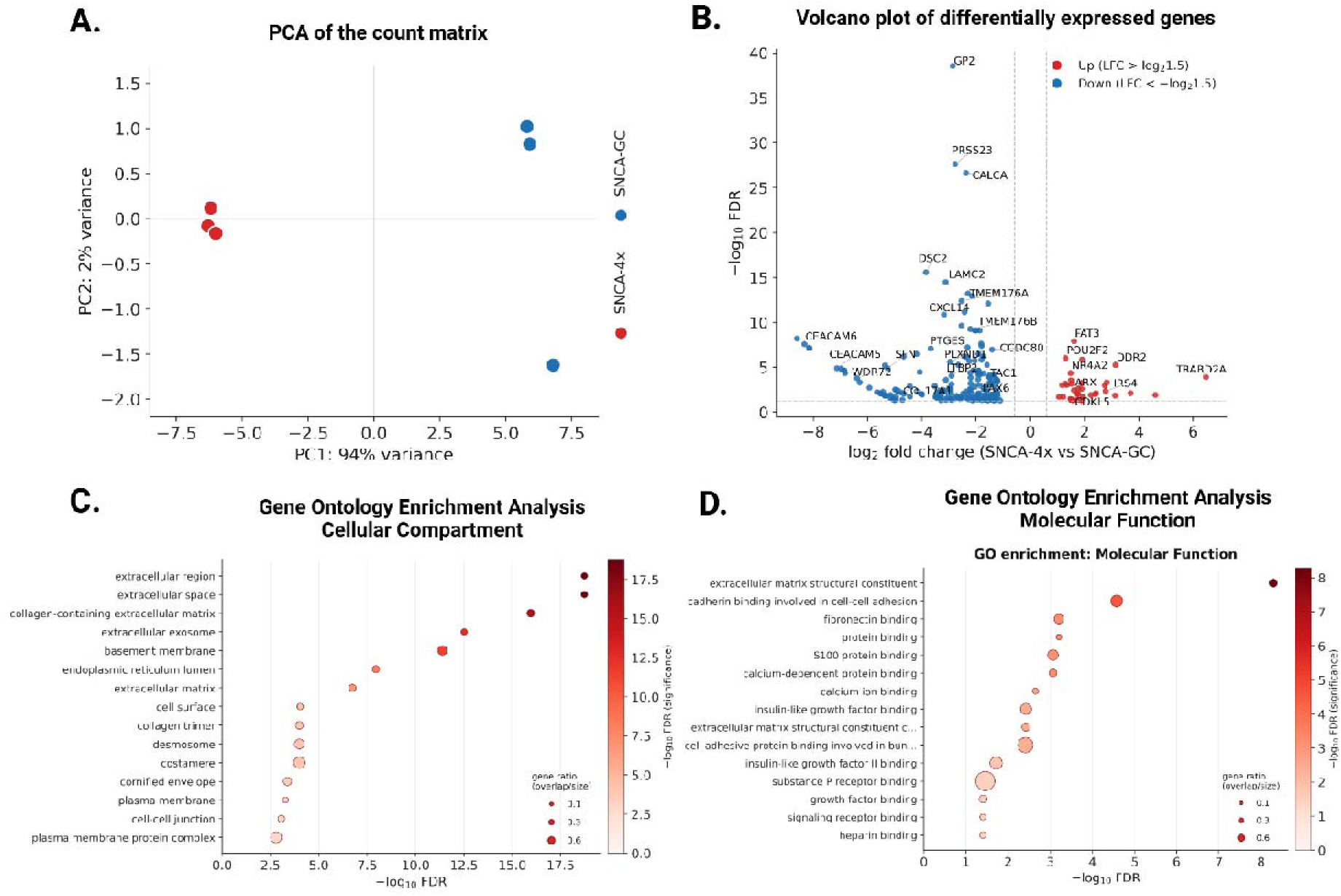
Differential gene expression analysis of iPSC-derived SNCA-4x and SNCA-GC neurons. (A) Principal component analysis (PCA) of iPSC-derived SNCA-4x mDANs (red) and SNCA-GC mDANs (blue), based on gene expression profiles from three biological replicates. (B) Volcano plot showing differentially expressed genes. Red dots indicate significantly downregulated genes, and blue dots indicate significantly upregulated genes in iPSC-derived SNCA-4x mDANs, compared to SNCA-GC mDANs. (C) Bubble plot of Gene Ontology enrichment analysis for cellular components, showing the top 15 altered cellular compartments in iPSC-derived SNCA-4x mDANs. (D) Bubble plot of Gene Ontology enrichment analysis for molecular function, showing the top 15 altered molecular functions in iPSC-derived SNCA-4x mDANs. Significantly enriched GO terms are shown after Benjamini-Hochberg false discovery rate (FDR) correction.

### 3.3 Establishment and validation of a neuronal EV isolation protocol

Given the transcriptional alterations in EV-related genes and pathways, we developed an optimized isolation protocol combining ultrafiltration, ultracentrifugation, and EV precipitation to obtain nEVs (Fig. 3A). Nanoparticle tracking analysis (NTA) confirmed that isolated vesicles ranged from 50 to 180 nm in diameter, consistent with small EV characteristics (Fig. 3B). Transmission electron microscopy (TEM) further validated the presence of intact EVs with characteristic morphology (Fig. 3C). Western blot analysis demonstrated enrichment of canonical EV markers, including the tetraspanins CD9 and CD63 and flotillin-1, while the endoplasmic reticulum marker calnexin was absent, confirming minimal cellular contamination (Fig. 3D). Importantly, the isolated vesicles were also enriched in NCAM, a neuronal surface marker (Fig. 3D), supporting their neuronal origin. Since our differentiation protocol yields cultures highly enriched for midbrain dopaminergic neurons, the resulting preparations are correspondingly highly enriched in neuron-derived EVs. We nevertheless cannot formally exclude a minor contribution from residual non-neuronal cells present in cultures, and therefore refer to these preparations as neuron-enriched rather than exclusively neuronal.

**Figure 3.**
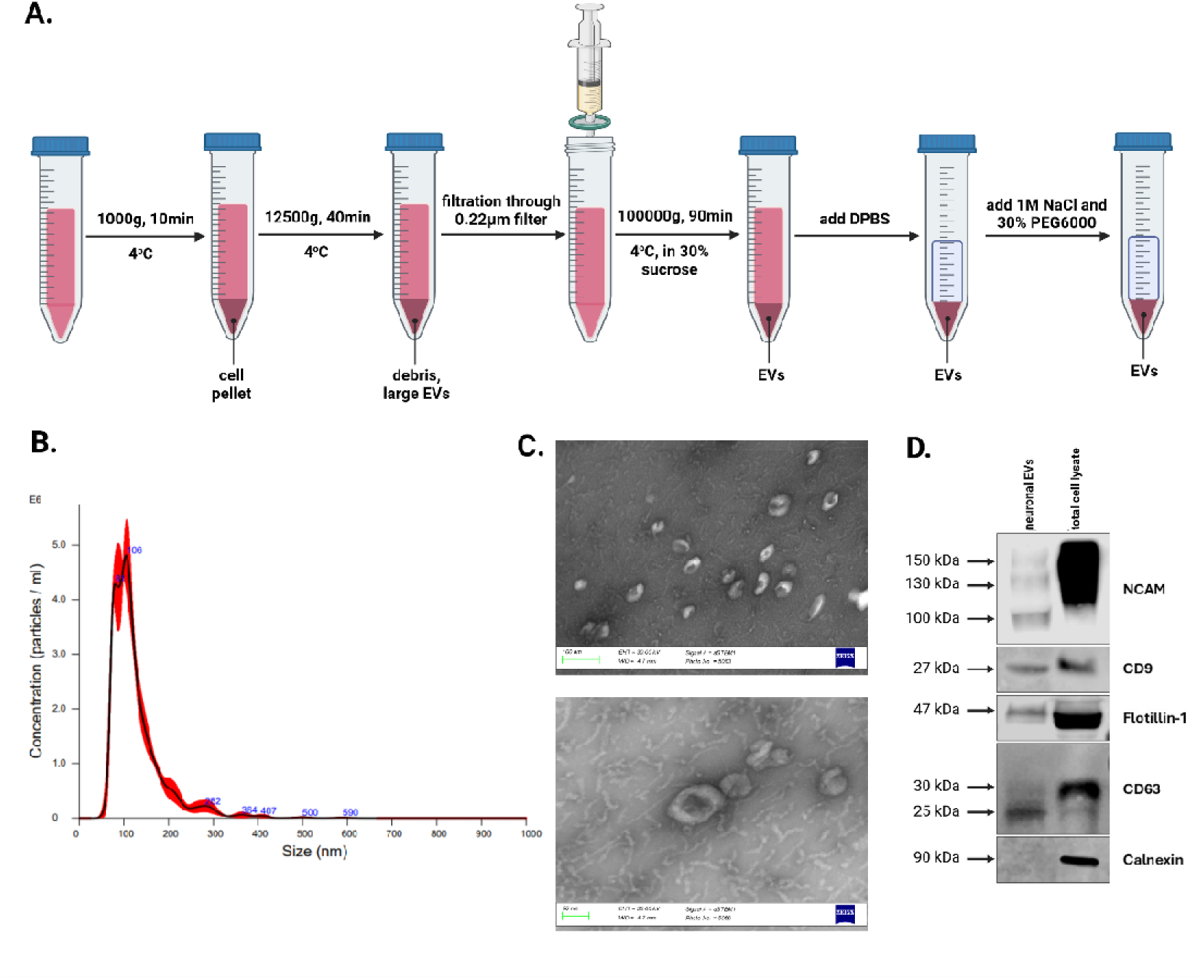
EV isolation and characterization. (A) Schematic representation of the extracellular vesicle isolation workflow, which combined ultrafiltration, ultracentrifugation, and polyethylene glycol (PEG) precipitation. (B) Nanoparticle tracking analysis (NTA) confirming the isolation of extracellular vesicles with an average diameter ranging from 50 to 180 nm. (C) Transmission electron microscopy (TEM) images confirming the isolation of extracellular vesicles. Scale bars: 100 nm (top) and 50 nm (bottom). (D) Western blot analysis showing that isolated nEVs are positive for the tetraspanins CD9 and CD63, as well as Flotillin-1 and NCAM, and negative for calnexin, in contrast to total cell lysate (TCL).

### 3.4 SNCA-4x mDANs secrete increased numbers of smaller EVs with altered molecular cargo

Using our validated isolation protocol, we investigated whether the increased α-synuclein levels of the SNCA-4x mDANs affect nEV secretion and composition. NTA revealed that SNCA-4x mDANs secreted significantly more nEVs compared to isogenic controls, with a modest but significant reduction in mean particle diameter (Fig. 4A). Quantification of α-synuclein content using a sensitive in-house ELISA demonstrated significantly elevated α-synuclein levels in SNCA-4x-derived nEVs compared to isogenic control-derived nEVs, while SNCA KO-derived neuronal EVs showed minimal detection signal, confirming assay specificity (Fig. 4B).

**Figure 4.**
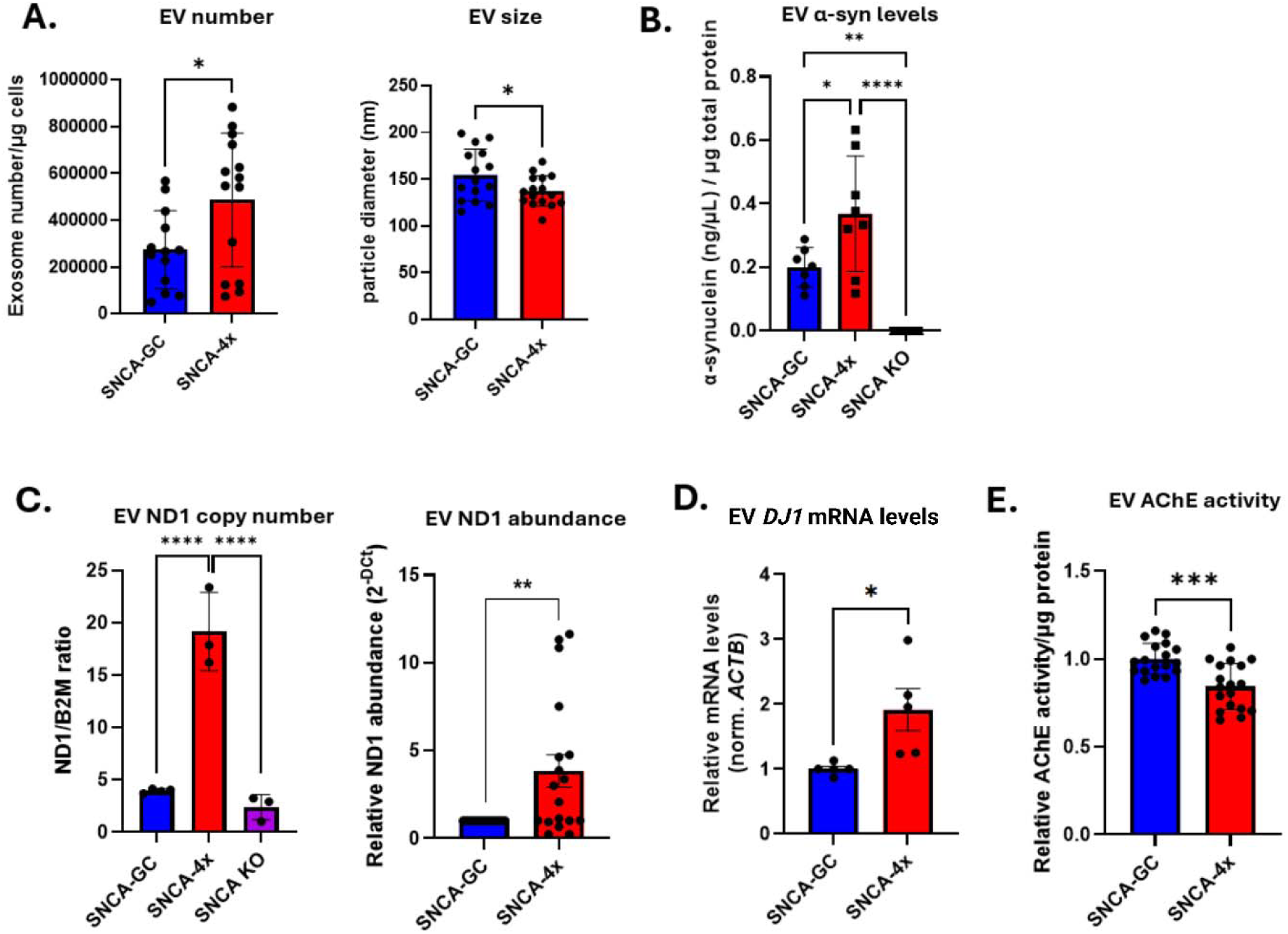
Differential EV features and cargo in iPSC-derived SNCA-4x and SNCA-GC neuronal EVs. (A) Differences in EV number and size between SNCA-4x-derived nEVs (red) and SNCA-GC-derived nEVs (blue), measured by NTA in six biological replicates. (B) Differences in α-synuclein protein levels in SNCA-4x-derived nEVs (red), SNCA-GC-derived nEVs (blue), and SNCA-KO-derived nEVs (purple), measured by ELISA in three biological replicates. (C) Differences in mtDNA levels in SNCA-4x-derived nEVs (red), SNCA-GC-derived nEVs (blue), and SNCA-KO-derived nEVs (purple), measured by digital PCR in four biological replicates, followed by differences in mtDNA levels between SNCA-4x-derived nEVs (red) and SNCA-GC-derived nEVs (blue), measured by qPCR for the ND1 gene in three biological replicates. (D) Differences in PARK7 mRNA levels between SNCA-4x-derived nEVs (red) and SNCA-GC-derived nEVs (blue), measured in three biological replicates. (E) Differences in AChE activity between SNCA-4x-derived nEVs (red) and SNCA-GC-derived nEVs (blue), measured in three biological replicates. For panels A, C2, D, and E, an unpaired t-test was performed. For panels B and C1, one-way ANOVA followed by Tukey’s multiple-comparison test was used. ns, not significant; \**p* < 0.05, \*\**p* < 0.01, \*\*\**p* < 0.001.

Given our previous findings that SNCA-4x mDANs exhibit mitochondrial dysfunction characterized by elevated reactive oxygen species and decreased mitochondrial membrane potential (Gorgogietas et. al., 2025), we evaluated whether mitochondrial components were differentially packaged into nEVs. Digital and quantitative PCR analyses revealed significantly increased mtDNA copy number in SNCA-4x-derived nEVs compared to isogenic control-derived nEVs (Fig. 4C). Interestingly, mtDNA levels appeared to correlate with α-synuclein expression, as SNCA KO-derived nEVs contained the lowest mtDNA copy numbers, followed by isogenic controls and SNCA-4x cells (Fig. 4C).

To further characterize nEV cargo, we quantified mRNA levels of PD-associated genes documented in EVs from the ExoRBase 2.0 database. Among the transcripts evaluated, we observed a significant increase in *PARK7* gene (encoding DJ-1) mRNA levels in SNCA-4x-derived nEVs compared to controls (Fig. 4D). Finally, based on reports that plasma EVs from PD patients exhibit decreased acetylcholinesterase (AChE) activity that negatively correlates with disease severity (Shim et al., 2021), we measured AChE enzymatic activity in nEVs. Consistent with patient data, SNCA-4x-derived nEVs showed modest but significantly reduced AChE activity compared to isogenic control-derived nEVs (Fig. 4E).

Collectively, these findings demonstrate that *SNCA* triplication alters the size, particle number and molecular cargo of secreted nEVs, affecting their protein, DNA, and RNA content. These results indicate that nEV composition correlates with neuronal dysfunction and support the value of nEVs for both mechanistic studies and biomarker development in PD research.

### 3.5 SNCA-4x-derived nEVs exhibit enhanced microglial uptake and trigger pro-inflammatory activation

To assess the functional impact of the altered nEV cargo described above, we exposed iPSC-derived microglia from the isogenic control line to nEVs derived from either SNCA-4x or SNCA-GC neurons. Untreated microglia and LPS+IFNγ-treated microglia were included as baseline and positive controls, respectively. We first evaluated vesicle uptake and found that microglia internalized SNCA-4x-derived nEVs approximately twice as efficiently as SNCA-GC nEVs (Fig. 5A-B). Furthermore, quantitative morphometric analysis revealed significant differences in microglial activation state, since microglia that had phagocytosed SNCA-4x nEVs showed increased circularity and solidity compared to microglia that phagocytosed control nEVs (Fig. 5C).

**Figure 5.**
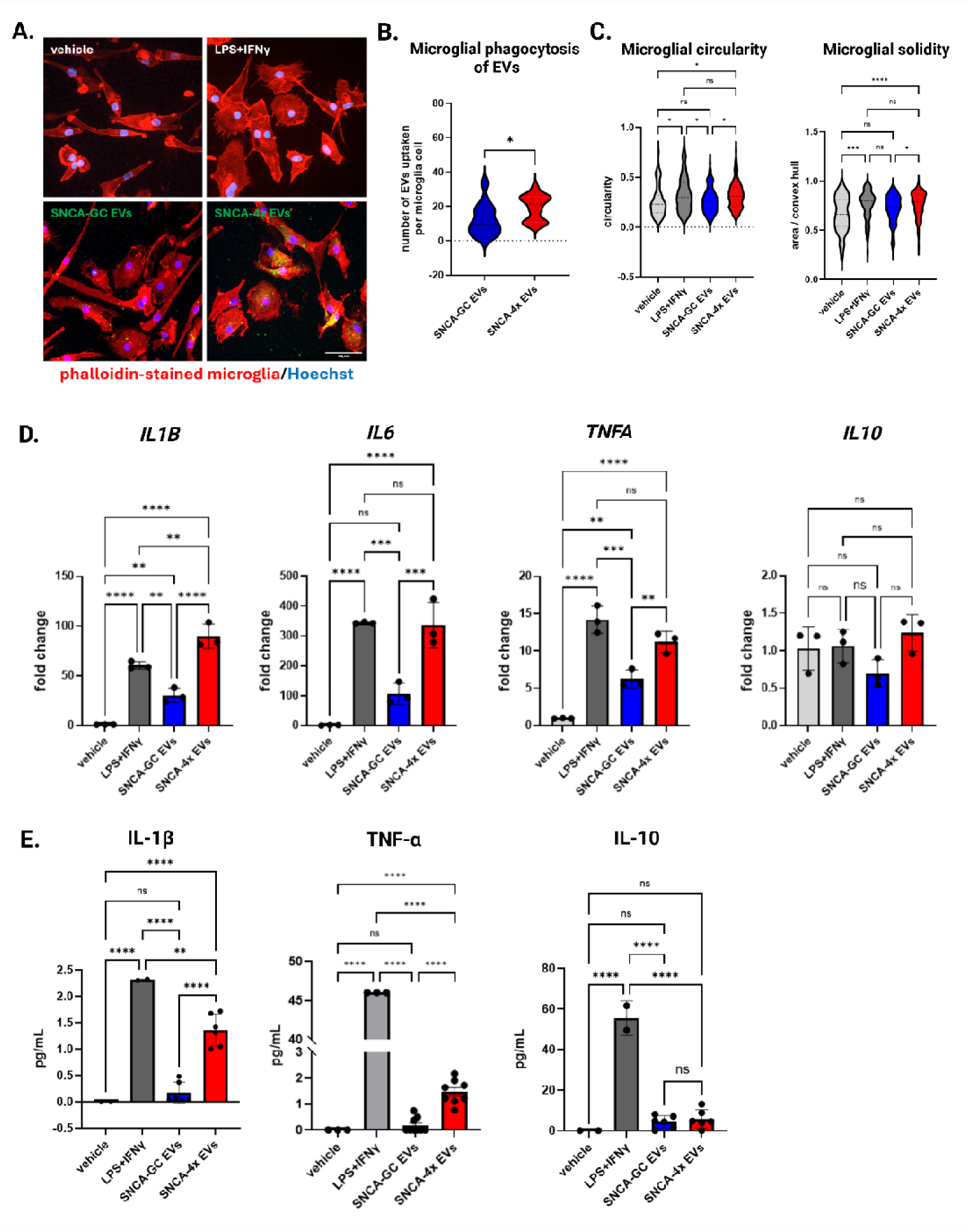
Neuronal EV uptake by SNCA-GC microglia and microglial response. (A) Representative fluorescence images of microglia stained with phalloidin (red) under the following conditions: untreated, treated with LPS + IFN-γ, treated with CFSE-labeled SNCA-GC nEVs (green), or treated with CFSE-labeled SNCA-4x nEVs (green). Nuclei were stained with Hoechst 33342 (blue). Scale bar: 50 μm. (B) Quantification of microglial uptake of SNCA-GC- or SNCA-4x-derived nEVs. (C) Quantification of microglial morphological features, including circularity and solidity, after treatment with LPS + IFN-γ, SNCA-GC EVs, SNCA-4x EVs, or under untreated conditions. (D) Relative mRNA expression of selected cytokines (IL-1β, IL-6, TNF-α, IL-10) in microglia after treatment with LPS + IFN-γ, SNCA-GC nEVs, SNCA-4x EVs, or under untreated conditions. (E) Absolute quantification of the protein levels of selected cytokines (IL-1β, TNF-α and IL-10) in microglial culture medium after treatment with LPS + IFN-y, SNCA-GC EVs, SNCA-4x EVs, or under untreated conditions. All experiments were performed in triplicate. For panel B, an unpaired t-test was used. For panels C–E, one-way ANOVA followed by Tukey’s multiple-comparison test was used. ns, not significant; \**p* < 0.05, \*\**p* < 0.01, \*\*\**p* < 0.001, \*\*\*\**p* < 0.0001.

To further assess whether nEV exposure initiates an inflammatory response, we quantified the mRNA expression levels of key cytokines. Because the recipient microglia and the nEV-donor neurons were differentiated from the same parental iPSC background, this setup constitutes an autologous neuron–microglia system in which donor genetic background is matched and SNCA copy number is the only variable. We observed a significant upregulation of *TNF-α, IL-6, IL-1 β*, and *IL-10* transcripts following exposure of microglia to SNCA-4x-derived EVs compared to both untreated controls and SNCA-GC EV-treated conditions. Notably, the magnitude of this increase was comparable to that induced by LPS + IFN-γ treatment. To validate these findings at the protein level, we subsequently quantified TNF-α, IL-1β, and IL-10. At the protein level, IL-10 was not altered under any condition. However, consistent with the transcriptional data, TNF-a and IL-1(3 levels were significantly elevated following SNCA-4x nEV exposure relative to both untreated controls and SNCA-GC nEV treatment.

## 4. Discussion

### 4.1 Principal findings

In this study, we demonstrate that iPSC-derived midbrain dopaminergic neurons harboring an *SNCA* triplication not only recapitulate key PD-related features such as accumulation of α-synuclein multimers and mitochondrial dysfunction (as shown by Gorgogietas et al., 2025), but also show profound alterations in the biogenesis, secretion, and cargo composition of neuronal extracellular vesicles. Bulk RNA-seq revealed that SNCA-4x mDANs display transcriptional changes in pathways related to EVs, membrane organization, and extracellular matrix dynamics, suggesting a re-wiring of vesicular trafficking and intercellular communication. Using a multi-step isolation protocol, we showed that SNCA-4x neurons release more EVs, which are on average smaller in size and enriched in α-synuclein, mtDNA, and *PARK7* mRNA, while their acetylcholinesterase (AChE) activity is significantly reduced. Furthermore, we demonstrate that SNCA-4x-derived nEVs exhibit enhanced uptake by iPSC-derived microglia compared to isogenic control-derived nEVs, triggering a robust pro-inflammatory response characterized by a significant upregulation of cytokine transcripts, with corresponding elevations in TNF-α and IL-1β protein levels. This inflammatory response indicates that SNCA-4x nEVs may act as potent drivers of microglial activation. These findings identify nEVs as sensitive indicators; their molecular composition correlates with neuronal dysfunction in a-synuclein-driven PD, and highlights specific features, such as α-synuclein protein levels, mtDNA, *PARK7* mRNA, and AChE activity, that are associated with the enhanced microglial activation caused by SNCA-4x nEVs, although we cannot exclude that additional, unmeasured cargoes contribute to this inflammatory phenotype. Together, these features represent promising candidate biomarkers that need further mechanistic investigation.

### 4.2 Comparison with prior studies

Our observation that *SNCA* triplication neurons secrete more extracellular vesicles is consistent with the broader concept that α-synuclein-driven proteostatic stress promotes EV-mediated disposal of misfolded proteins and organelles. Elevated α-synuclein caused by *SNCA* triplication has been shown to impair neuronal differentiation and increase autophagic flux, indicating that proteostatic stress is already present well before overt neurodegeneration (Oliveira et al., 2015; Serra-Almeida et al., 2026). In *SNCA* triplication human midbrain organoids, transcriptomic and proteomic analyses revealed early changes in proteostasis-related signaling and extracellular matrix organization, suggesting that *SNCA* overexpression broadly reprograms neuronal vesicular and intercellular communication pathways before cell loss (Statoulla et al., 2026; Patikas et al., 2023). In parallel, oxidative stress and α-synuclein aggregation have been shown to trigger the release of EVs containing α-synuclein oligomers, which can be internalized by recipient neurons and compromise their survival. This evidence derives largely from immortalized human lines with artificial α-synuclein overexpression (H4 neuroglioma, Danzer et al., 2012 and inducible SH-SY5Y, Emmanouilidou et al., 2010) and from rodent primary neurons combined with overexpressing cell lines (Zhang et al., 2018). By contrast, EV cargo and function have not, to our knowledge, been characterized in isogenic human iPSC-derived midbrain dopaminergic neurons in which α-synuclein dosage is the only variable, where nEV alterations arise from endogenous *SNCA* copy number on a genetically matched background rather than from transgene overexpression. These data collectively support the idea that SNCA-related proteostatic stress promotes EV secretion as a compensatory or stress-induced mechanism, aligning with our finding of increased nEV release from SNCA-4x neurons.

In line with this, our data show enrichment of monomeric α-synuclein in nEVs, which is predominantly presented on the vesicle surface, mirroring experimental work on EV-associated α-synuclein. Using cultured primary neurons of mice models expressing A53T α-synuclein, Zhang et al. demonstrated that EVs contain α-synuclein oligomers and that the protein is located both inside the vesicles and on their surface, where it facilitates EV internalization by recipient neurons (Zhang et al., 2018). In those studies, EV-associated multimeric α-synuclein was shown to be internalized by axons of healthy neurons, leading to degeneration, and the vesicular α-synuclein was found to be partially accessible for protease digestion, confirming surface exposure (Zhang et al., 2018). In our system, a proteinase K protection assay confirmed that α-synuclein was not resistant to proteinase K treatment alone and was lost upon co-treatment with Triton X-100 (Figure S3A), indicating that the bulk of nEV α-synuclein is not mainly enclosed within the vesicle lumen but rather exposed on the surface. Mechanistically, α-synuclein’s strong affinity for negatively charged phospholipids and membrane biomolecules has been linked to its recruitment to lipid raft microdomains and to vesicular membranes during EV biogenesis (Ugalde et al., 2019; Gustafsson et al., 2018; Pirc & Ulrih, 2015), which is consistent with the idea that α-synuclein is enriched on the EV surface in our system. Our ELISA-based measurements capture predominantly monomeric EV α-synuclein, but the existing literature supports a model in which EV-associated α-synuclein includes both monomeric and oligomeric species, capable of seeding pathology and modulating intercellular communication (Gustafsson et al., 2018; Zhang et al.,2018; Chistiakov & Chistiakov, 2017).

Our finding that SNCA-4x-derived nEVs carry elevated mtDNA content is consistent with the concept that mitochondrial stress promotes packaging of mitochondrial components into EVs. Studies of EVs released under oxidative stress or α-synuclein-rich conditions have shown that EVs can contain mitochondrial proteins and mtDNA and can transmit these to recipient cells, with opposite consequences depending on cargo quality. Transfer of intact mitochondria can be supportive, as astrocyte-derived mitochondrial particles entering neurons after ischaemia amplify survival signalling and blocking this transfer worsens outcome (Hayakawa et al., 2016), whereas oxidized components and free mtDNA instead act as DAMPs. Cells normally gate this distinction, excluding oxidized, pro-inflammatory material from EVs via a Parkin-dependent route to lysosomes, so that cargo quality rather than quantity determines the paracrine effect (Todkar et al., 2021; Manzar et al., 2025). Moreover, a growing body of evidence indicates that mitochondrial-derived vesicles or mito-EVs can serve as carriers of mitochondrial DAMPs, thereby contributing to neuroinflammation and disease progression in CNS disorders (Deus et al.,2022; Nakamya et al., 2022; Kim et al., 2024). In PD and related synucleinopathies, EV-mediated transmission of monomeric and multimeric α-synuclein and mitochondrial cargo has been proposed as a mechanism that links neuronal stress to glial activation and chronic neuroinflammation (Deus et al., 2022; Guo et al., 2020; Nakamya et al., 2022). In previous work using the same SNCA-4x iPSC-derived midbrain dopaminergic neurons, we showed that a-synuclein accumulation is accompanied by mitochondrial dysfunction, characterized by elevated reactive oxygen species and reduced mitochondrial membrane potential relative to isogenic gene-corrected controls, as determined by high-content morphological profiling of the neurons themselves (Gorgogietas et al., 2025). However, that study did not assess extracellular vesicles. The elevated mtDNA we now detect in SNCA-4x-derived nEVs therefore extends this intracellular mitochondrial phenotype to the vesicular compartment and is consistent with reports that mitochondrial stress promotes the release of mtDNA-carrying EVs (Deus et al., 2022; Zhou et al., 2023). Whether the mitochondrial dysfunction previously described is causally upstream of the increased nEV mtDNA load remains to be established.

The increase in *PARK7* mRNA in SNCA-4x-derived nEVs is consistent with a role for DJ-1 in the neuronal response to α-synuclein-driven oxidative stress. DJ-1 is a redox-sensitive protein implicated in autosomal recessive PD, and its deficiency sensitizes embryonic stem cell-derived dopamine neurons to oxidative stress and proteasomal inhibition (Martinat et al., 2004), while DJ-1 is upregulated and redistributed following oxidative insult (Lev et al., 2008). Critically for our model, DJ-1 also acts as a redox-dependent chaperone that suppresses α-synuclein aggregation (Shendelman et al., 2004). Its induction in neurons carrying four *SNCA* copies is therefore mechanistically coherent, engaging a system directly opposed to the protein these cells overproduce. Direct evidence that DJ-1 is EV-associated in humans comes from patient plasma rather than from cellular models. Using L1CAM immunocapture to enrich neuron-derived exosomes from the plasma of PD patients and controls, Zhao and colleagues found DJ-1 and a-synuclein protein elevated in the neuron-derived fraction, together with an increased ratio of exosomal to total DJ-1, whereas unfractionated plasma levels did not differ between groups (Zhao et al., 2018). That study was descriptive, quantifying protein by ELISA without testing whether EV-associated DJ-1 influences recipient cells, and no association with disease progression was detected. Notably, the identification of neuron-derived EVs in blood remains methodologically contested, as L1CAM may circulate largely as free protein (Norman et al., 2021), although single-vesicle analyses subsequently confirmed L1CAM-positive nEVs in blood (Nogueras-Ortiz et al., 2024); our nEVs avoid this ambiguity, being neuronal by origin rather than by marker capture. Our data therefore extend this concept by showing that PARK7 mRNA is enriched in EVs from SNCA-4x neurons, which may represent a stress-associated EV cargo whose biological significance requires future investigation. Beyond PARK7, we found that the integrated stress-response transcript *ATF4* was enriched in SNCA-4x-derived nEVs relative to isogenic controls, with a similar trend for *PINK1*, even though their cellular levels did not differ between genotypes (Figure S2). This observation supports selective sorting of stress-response transcripts into nEVs rather than passive leakage, reinforcing the application of nEV cargo as an active readout of neuronal stress.

Additionally, our finding of reduced AChE activity in SNCA-4x nEVs relates to clinical studies of EV AChE in PD, although the comparison requires careful consideration of both the biomaterial and the vesicle subpopulation examined. In bulk EV isolated by ultracentrifugation from the plasma of PD patients (a preparation of mixed cellular origin, with no cell-type selection), AChE activity was significantly reduced relative to healthy controls and correlated negatively with disease severity (Hoehn & Yahr, UPDRS part III), suggesting that EV AChE may reflect peripheral cholinergic dysfunction and disease progression (Shim et al., 2021). A subsequent study fractionated plasma EVs by immunocapture using CD9, a tetraspanin present on vesicles from many cell types and therefore a general EV marker rather than a cell-type marker, and L1CAM, a neuronal adhesion molecule used to enrich neuron-derived EVs. AChE activity was markedly decreased in the CD9-positive, mixed-origin fraction from PD patients, whereas the L1CAM-positive neuronal fraction showed no significant difference (Jeong et al., 2024). The reduction detectable in patient plasma therefore localizes to the general, largely peripheral EV pool rather than to vesicles of neuronal origin, indicating a subpopulation-restricted rather than a universal change. Our data differ from these observations: we detect reduced AChE activity in vesicles of defined neuronal origin, released directly by iPSC-derived mDANs, whereas the only clinical fraction enriched for nEVs showed no change (Jeong et al., 2024). This might be explained by the starting biomaterial: we sample nEVs directly at their neuronal source, whereas in plasma these vesicles are diluted within a much larger pool of peripheral EVs. The reduction of AChE activity may also reflect lower AChE expression, post-translational modification, or a shift toward EV subpopulations carrying less active enzyme. Our data therefore extend rather than replicate the clinical findings and indicate that EV AChE is best assessed by vesicle subtype and cellular origin rather than as a generic EV measure.

Our data show that nEVs from SNCA-4x neurons are taken up approximately twice as efficiently by microglia as those from isogenic controls and drive a robust pro-inflammatory response, with increased *TNF-α, IL-6* and *IL-1fi* transcripts and elevated TNF-α and IL-1β protein. Because recipient microglia were genetically identical across conditions, this difference is due to the nEV cargo rather than to the responder cell. This is direct evidence that the direction of EV-mediated neuron–microglia signaling is set by what the donor neuron packages in EVs, which in turn reflects its physiological state. Peng and colleagues showed that EVs from healthy neurons act homeostatically, inhibiting microglial activation and reducing LPS-induced TNF-α, IL-6 and MCP-1 while increasing IL-10 (Peng et al., 2021). SNCA-4x nEVs, enriched in α-synuclein and mtDNA (both established DAMPs), invert this outcome, indicating that the same vesicular route switches from homeostatic to pro-inflammatory, when the donor neuron is under proteostatic and mitochondrial stress.

This pro-inflammatory effect of PD-associated EVs is consistent with multiple studies demonstrating that α-synuclein-carrying EVs act as “deadly transmitters” (Chistiakov & Chistiakov, 2017) that propagate pathology and inflammation. Neuron-derived EVs were found to transfer monomeric and multimeric α-synuclein forms between neuronal and non-neuronal cells (Emmanouilidou et al.,2010; Alvarez-Erviti et al., 2011a; Zhang et al., 2018; Mavroeidi et al., 2022), including microglia, thereby contributing to α-synuclein spreading, and deposition of α-synuclein in glial cells induces inflammation that propagates to other glial cells and neurons. Extracellular vesicles derived from the plasma of PD patients (PD-EVs) have recently been shown to drive microglial senescence and neurodegeneration, inducing a proinflammatory, senescence-associated secretory phenotype (SASP) with elevated TNF-α, IL-1β, IFN-γ, IL-8, and CCL11 in HMC3 microglial cells, and proteomic analysis revealed enrichment of proteins involved in complement cascades and immune response (Yadav et al., 2026, Preprint). These findings require caution, as HMC3 cells are SV40 large T–immortalized and SV40 large T inactivates p53 and Rb, the principal effectors of senescence, making this line poorly suited to studying SASP induction. By contrast, our non-transformed iPSC-derived microglia express *TMEM119* and *P2RY12* (Fig. 1E) and are genetically matched to the neurons supplying the nEVs, which are of defined neuronal rather than mixed plasma origin. Furthermore, α-synuclein itself acts as a potent inflammatory stimulator of microglial cells, inducing large increases in p38 MAP kinase, ERK1/2, and JNK pathways within minutes, with secretions containing increased IL-1[3 and TNF-α that are toxic to neuronal cells (Correa & Eales, 2012; Klegeris et al., 2008; Wang et al., 2023).

The elevated mtDNA in SNCA-4x-derived nEVs further supports the mechanism of inflammation induction, as mtDNA is a well-characterized DAMP that activates microglia. Mitochondrial-derived vesicles (MDVs) enriched in mtDAMPs can induce immune responses and release of pro-inflammatory cytokines, with mtDNA sensed by microglial cells through pathways promoting proinflammatory phenotype perpetuation via NLRP3 inflammasome activation (Deus et al., 2022; Qiu et al., 2022; West et al., 2015; Wasner et al., 2022). Microglial EVs have also been shown to contribute to α-synuclein pathology progression by altering intracellular α-synuclein accumulation and increasing protein clearance dysregulation, using primary healthy microglia and BV2 models, suggesting bidirectional EV-mediated crosstalk between neurons and microglia in PD (Xia et al., 2019; Guo et al., 2020; Turola et al., 2012). Additionally, total plasma EVs from PD patients contain significantly increased pro-IL-1 β and TNF-α levels compared with controls, supporting the concept of inflammatory pathogenesis in PD development and progression (Chan et al., 2021; Yadav et al., 2026, Preprint).

### 4.3 Strengths and limitations

A major strength of this study is the isogenic design, which allows direct comparison of *SNCA* triplication (SNCA-4x), gene-corrected control (SNCA-GC), and SNCA knockout (SNCA-KO) lines while minimizing genetic background variability. This extends to the recipient microglia, which were deliberately derived from the control line: because microglia express substantially lower endogenous α-synuclein than neurons, a control-genotype responder differs from the nEV-donor neurons only in SNCA copy number. This avoids adding microglial pathology on top of the nEV effect, so the response we measured reflects the nEV cargo alone, and makes the finding more general, since it does not depend on the recipient cells’ own genotype. Furthermore, combining RNA-seq with a multi-step EV isolation protocol enabled robust characterization of vesicular properties and molecular cargo, including EV-specific markers and the absence of endoplasmic reticulum contamination. Moreover, the integration of multiple complementary readouts (ELISA, qRT-PCR, dPCR, and functional assays) provides a comprehensive view of protein, nucleic acid, and enzymatic alterations in EVs. The inclusion of iPSC-derived microglia for EV uptake and inflammation assays enables direct modeling of neuron–microglia communication in a human genetic PD system.

Nonetheless, several limitations should be acknowledged. First, our neuronal cultures are relatively simplified compared with the *in vivo* nigrostriatal circuit, lacking the full complexity of glial and vascular interactions. Second, while we used a defined differentiation protocol, subtle differences in maturation state or cell-type composition may influence EV profiles, even within isogenic lines. Third, our ELISA-based α-synuclein measurements report total EV α-synuclein without direct resolution of monomeric versus oligomeric forms, so we cannot definitively distinguish which species are enriched. Fourth, our current EV-uptake experiments do not distinguish between direct cargo-dependent inflammatory signaling (e.g., mtDNA via NLRP3, α-synuclein via surface receptors) versus indirect effects mediated by EV surface properties; mechanistic dissection of the specific DAMP-receptor interactions driving microglial activation requires future studies. Finally, our current EV-uptake experiments in microglia focus on morphological and early cytokine readouts; longer-term functional outcomes, such as sustained microglial activation or synaptic alterations, remain to be explored.

### 4.4 Biological and translational implications

Biologically, our results support a model in which α-synuclein overexpression and *SNCA* triplication drive neuronal stress that is partially compensated, or inadvertently propagated, by altered EV secretion. Increased release of smaller nEVs carrying higher levels of α-synuclein, mtDNA, and *PARK7* mRNA may serve as a mechanism for transmitting pathological signals to neighboring cells. The reduced AChE activity in nEVs further suggests that cholinergic-related pathways are perturbed even at the level of secreted vesicles, pointing toward broader synaptic and neuromodulatory defects beyond the dopaminergic system. In parallel, the enhanced uptake of SNCA-4x-derived EVs by iPSC-derived microglia implies that EV-mediated crosstalk may contribute to early neuroinflammatory responses, consistent with the concept that EVs act as carriers of DAMPs and proteinopathies in the brain.

From a translational perspective, our findings identify several EV-associated features that could be developed into biomarkers or therapeutic targets. The SNCA-dependent increases in α-synuclein, mtDNA, and *PARK7* mRNA in nEVs provide candidate molecular signatures that, once validated in patient-derived EVs, might be used for early detection or stratification. Similarly, the reduction in AChE activity in EVs parallels prior clinical observations and may offer a peripheral readout of cholinergic dysfunction that could supplement existing clinical scores. These findings also expose the advantages of an isogenic iPSC over the approaches that have dominated EV research in PD. Existing evidence derives from immortalized cell lines with artificial α-synuclein overexpression (H4 neuroglioma, Danzer et al., 2012; inducible SH-SY5Y, Emmanouilidou et al., 2010) or from transformed microglial lines that express α-synuclein at non-physiological levels and lack a genetically matched control. On the other hand, patients’ biofluids include EVs of mixed cellular origin in which neuron-derived vesicles are a minority or are enriched using immunocapture markers of contested specificity (Norman et al., 2021; Nogueras-Ortiz et al., 2024). Our system avoids both constraints: nEVs are neuronal by origin rather than by capture-marker, α-synuclein is expressed from its endogenous gene and the SNCA-4x, SNCA-GC and SNCA-KO group attributes EV phenotypes to gene dosage on a common genetic background. This lets us link the EV changes directly to *SNCA* expression, which neither approach allows, although our cultures lack the complexity of the intact brain. Overall, our work strengthens the rationale for using iPSC-derived nEVs as a platform to dissect disease-specific mechanisms and to prioritize biomarker panels for future validation in biofluids from PD patients.

## 5. Conclusions

This study demonstrates that α-synuclein overexpression reshapes the biogenesis, molecular composition, and functional impact of neuronal extracellular vesicles, linking α-synuclein-driven proteostatic and mitochondrial stress to altered intercellular communication. The enrichment of α-synuclein, mtDNA, and enhanced microglial activation positions nEVs as candidate mediators and sensitive indicators of PD– related pathology. These findings support the use of iPSC-derived EVs for dissecting disease mechanisms and prioritizing EV-based markers as candidates for translational validation.

## Acknowledgements

We thank **Dr. Elizabet Petkovski** (Laboratoire National de Santé, Luxembourg) for performing the Short Tandem Repeat (STR) analysis of our cell lines. We also thank **Dr. Tilo Kunath** (University of Edinburgh) for providing the AST23, AST23-2KO, and AST23-4KO iPSC lines. We are grateful for the constant discussions with all the members of the Translational Neuroscience group of the Luxembourg Centre for Systems Biomedicine (LCSB), which improved this work. Finally, we thank **Dr. Dionysios Antonopoulos** (University of Thessaly) for his crucial assistance regarding EV isolation and characterization approaches.

## Author contributions

**Ioanna Boumpoureka** performed experiments, conducted data analysis, and contributed to figure preparation. **Sylvie Delcambre, Felix B. Kleine Borgmann**, **Johanna Heider** and **Arnaud Poret** performed experiments and carried out data analysis. **Johannes H. Wilbertz, Peter Sommer, Ibrahim Boussaad, Michel Mittelbronn,** and **Anne Grünewald** contributed to the discussion of the results and critically reviewed the manuscript. **Rejko Krüger** supervised the study, contributed to the discussion of the results, critically reviewed the manuscript, and secured funding. **Vyron Gorgogietas** conceptualized the study, performed experiments, conducted data analysis, wrote the original draft of the manuscript, and secured funding. **Vyron Gorgogietas** and **Rejko Krüger** contributed equally to this work and are co-senior authors.

## Data availability

All raw data, processed datasets, and analyses are available from the corresponding author upon reasonable request, subject to LCSB data protection policies. The data supporting this study derive from patient iPSC lines and constitute personal data under the GDPR. The informed consent obtained does not authorize public deposition or secondary sharing; access is therefore restricted.

## Ethics approval

All the experimental procedures performed in the context of this study are harmonized with the Declaration of Helsinki (the latest version of June 1964-World Medical Association) and are ethically covered by the approval provided by the National Ethics Board of Luxembourg (Comité National d’Ethique dans la Recherche; CNER #201411/05) to Prof. Rejko Krüger.

