## Supplementary file for "*SNCA* triplication shapes neuronal extracellular vesicle biology and promotes microglial activation in patient-derived iPSC-based models of Parkinson’s disease"

**Materials and Methods**

**S1. Neurofilament light chain (NF-L) ELISA**

Neurofilament light chain (NF-L) was quantified in nEV lysates using the Human NF-L ELISA Kit (Proteintech, Cat. No. KE00305), a sandwich ELISA, according to the manufacturer's instructions. Briefly, 50 μL of nEV lysates were diluted in the provided sample diluent and loaded, together with the kit standards, onto the pre-coated 96-well plate, and incubated to allow antigen capture. After washing, wells were incubated with the biotinylated anti-NF-L detection antibody, washed again, and incubated with HRP-conjugated streptavidin. Following a final wash, the reaction was developed with TMB substrate, stopped with the provided stop solution, and the absorbance was measured at 450 nm on a Cytation 5 plate reader (BioTek). NF-L concentrations were interpolated from a standard curve generated with the recombinant NF-L standards supplied with the kit, and values were normalized to nEV total protein.

**S2. Proteinase K / Triton X-100 protection assay**

To determine whether nEV-associated proteins were luminal, and therefore protected by an intact vesicle membrane, or exposed on the vesicle surface, isolated 50 μg of nEVs were subjected to a proteinase K protection assay. Equal aliquots of SNCA-4x or SNCA-GC-derived nEVs resuspended in 1X DPBS were divided into three conditions: (i) DPBS alone (untreated control), (ii) 100 ng/ml proteinase K in the presence of 1% Triton X-100, and (iii) 100 ng/ml proteinase K . Samples were incubated for 1 h at 37°C. In this design, surface-exposed proteins are accessible to proteinase K and are digested, luminal proteins are shielded by the intact membrane and remain protected, and permeabilization of the membrane with Triton X-100 renders luminal proteins accessible to digestion. After incubation, the reaction was stopped by the addition of SDS-PAGE loading buffer and heating at 95°C for 5 min, and samples were analysed by Western blotting as described in Section 2.3. The membrane marker profile and the EV cargo of interest, including α-synuclein and DJ-1, were compared across the three conditions to distinguish surface-associated from luminal localization.

**Tables**

**Table S1.**

| **Gene** | **Forward primer** | **Reverse primer** |
| --- | --- | --- |
| *ATF4* | 5’-TTCTCCAGCGACAAGGCTAAGG-3’ | 5’-CTCCAACATCCAATCTGTCCCG-3’ |
| *PINK1* | 5’-CAAGAGGCTCAGCTACCTGCAC-3’ | 5’-TGTCTCACGTCTGGAGGCACT-3’ |

**Figure legends**

**Figure S1. Cellular and neuronal EV mitochondrial DNA integrity, transcription, and copy number** **across genotypes**. Digital PCR quantification of mitochondrial DNA (mtDNA) parameters in SNCA-GC (blue), SNCA-4x (red), and SNCA-KO (grey) samples. (A) Cellular mtDNA deletion load (ND4/ND1 ratio). (B) Cellular mtDNA transcription (D-loop/ND1 ratio). (C) Cellular mtDNA copy number per cell (ND1/B2M ratio). (D) Neuronal EV mtDNA deletion load (ND4/ND1 ratio). (E) Neuronal EV mtDNA transcription (D-loop/ND1 ratio). Data are shown for three to four biological replicates. For panels A, B, and C, one-way ANOVA followed by Tukey's multiple-comparison test was used; for panels D and E, one-way ANOVA followed by Tukey's multiple-comparison test was used. ns, not significant; *p < 0.05, **p < 0.01.

**Figure S2. Stress-response and DJ-1 transcript levels in cells and neuronal EVs.**
Relative mRNA levels normalized to *ACTB* in SNCA-GC (blue), SNCA-4x (red), and SNCA-KO (grey) samples. (A) Cellular *PINK1* mRNA. (B) Cellular *ATF4* mRNA. (C) Cellular *DJ1* (*PARK7*) mRNA. (D) Neuronal EV *PINK1* mRNA in SNCA-GC and SNCA-4x. (E) Neuronal EV *ATF4* mRNA in SNCA-GC and SNCA-4x. Data are shown for three biological replicates. For panels A, B, and C, one-way ANOVA followed by Tukey's multiple-comparison test was used; for panels D and E, an unpaired t-test was performed. ns, not significant; *p < 0.05.

**Figure S3. Surface localization of neuronal EV cargo and neuronal EV NF-L levels.**
(A) Proteinase K protection assay on SNCA-GC and SNCA-4x-derived neuronal EVs, analysed by Western blot for DJ-1 (21 kDa) and α-synuclein (15 kDa). Vesicles were left untreated, treated with 100 ng/ml proteinase K (ProtK) in the presence of 1% Triton X-100 (TX100), or treated with 100 ng/ml proteinase K alone, as indicated below the blot. Densitometric quantification of DJ-1 and α-synuclein across the treatment conditions is shown on the right. Data shown for one biological replicate. (B) NF-L concentration in SNCA-GC and SNCA-4x-derived neuronal EVs, measured by ELISA and expressed in ng/mL. Data are shown for three biological replicates. For panel B, an unpaired t-test was performed. *p < 0.05.

**Figures**

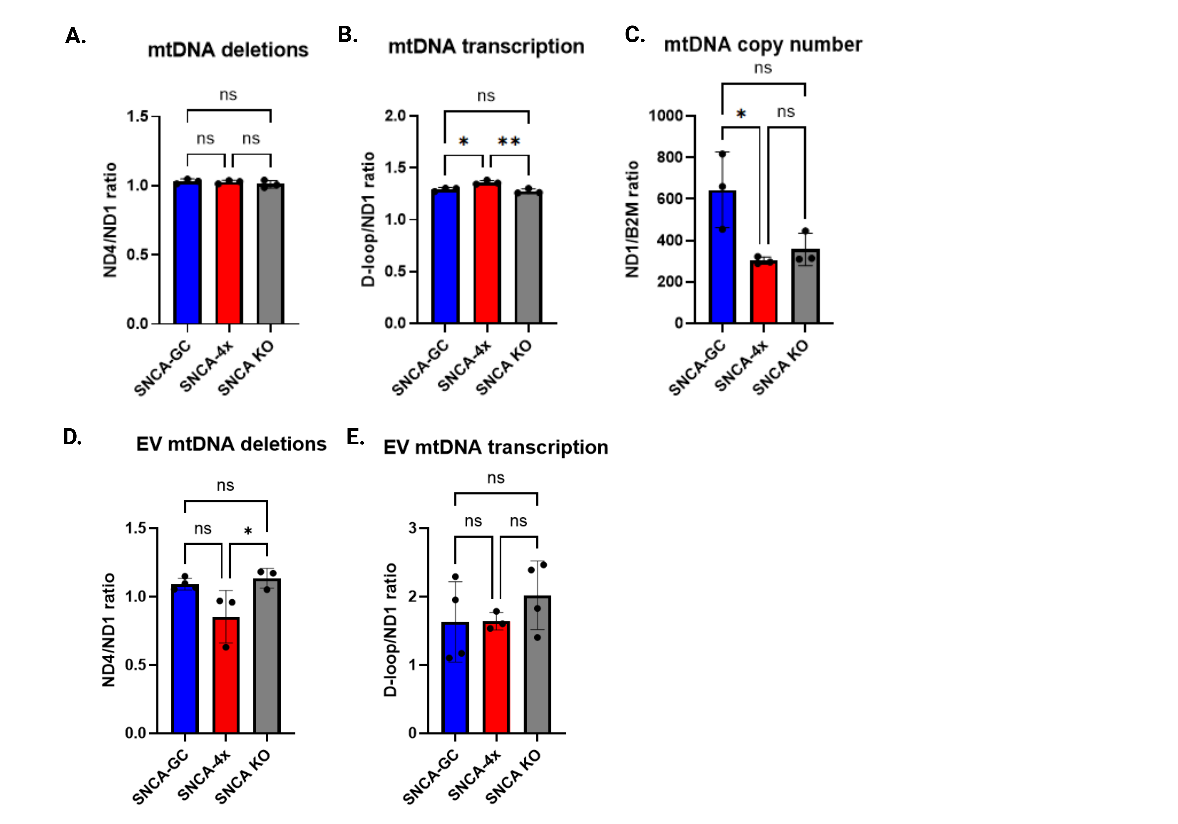
**Figure S1.**

**Figure 2.**

**Figure S2.**

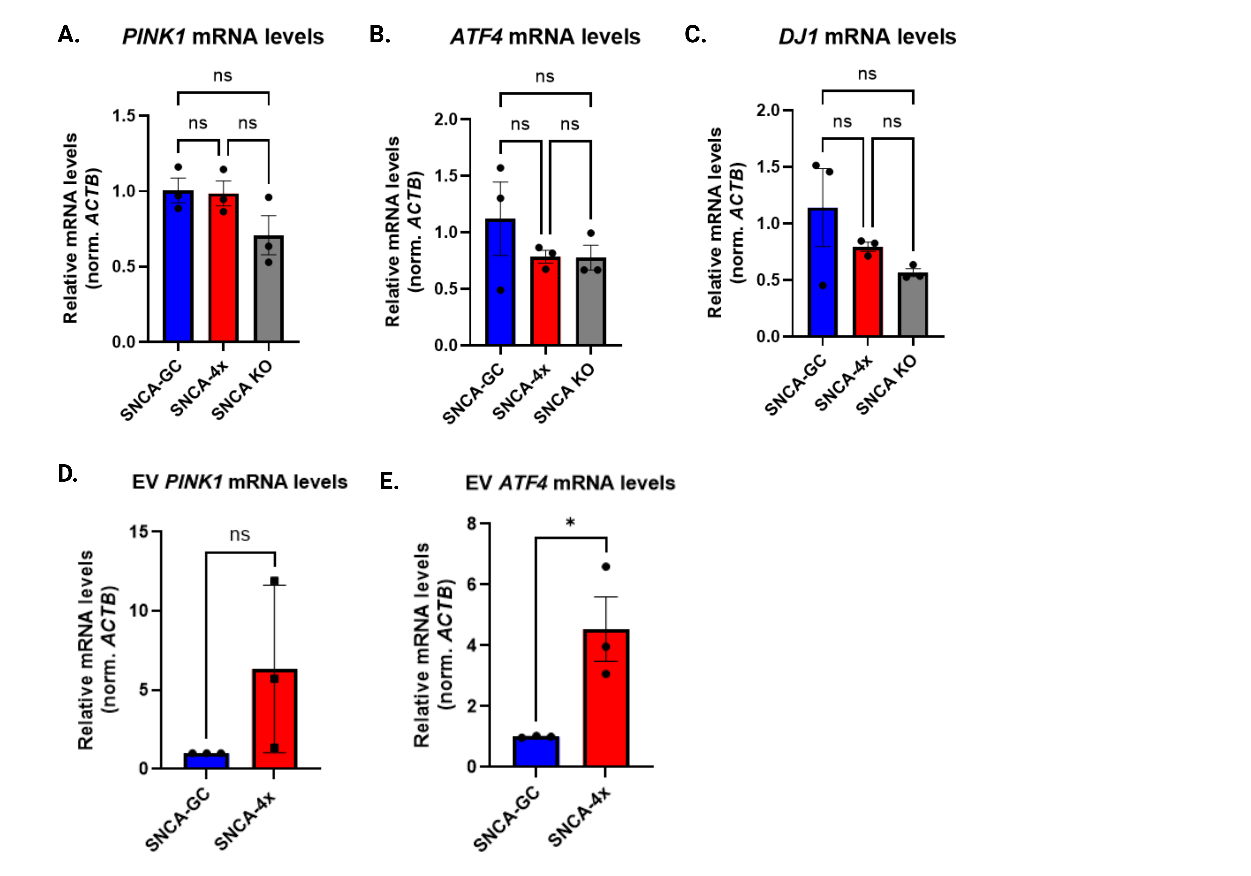

**Figure S3.**

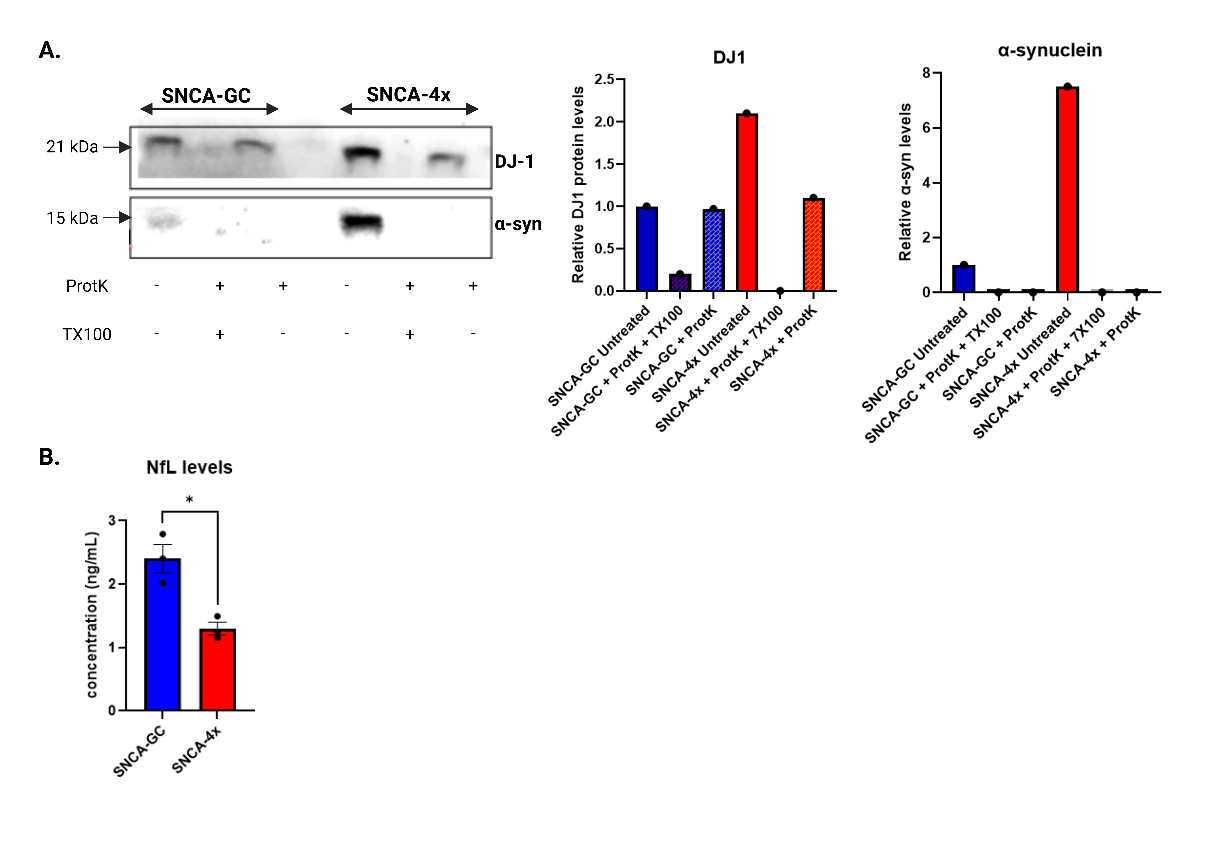
